# Mitochondrial transfer mediates metabolic communication between beta cells and islet macrophages

**DOI:** 10.64898/2026.09.01.748415

**Authors:** Lauar de Brito Monteiro, Anne-Sophie Archambault, Paula Ma, Galina Soukhatcheva, Derek Dai, Jane Velghe, Dorian Izerable, Majid Mojibian, Annette E. Patterson, Ramon I. Klein Geltink, C. Bruce Verchere

**Affiliations:** Department of Surgery, Faculty of Medicine, University of British Columbia, Canada; BC Children’s Hospital Research Institute, Vancouver, BC; Department of Pathology and Laboratory Medicine, Faculty of Medicine, University of British Columbia, Canada; Edwin S.H. Leong Centre of Healthy Aging, University of British Columbia, Vancouver, Canada; BC Cancer Research Centre, Department of Basic and Translational Research, Vancouver, BC, Canada; Centre for Molecular Medicine and Therapeutics, University of British Columbia, Vancouver, BC, Canada

## Abstract

Pancreatic islet macrophages support islet homeostasis and adapt their metabolic program in response to environmental cues, including beta cell released factors. Intercellular mitochondrial transfer is a biological process that modulates cellular responses. To test whether beta cells, which are strongly secretory, transfer mitochondria to islet macrophages, we generated mice with beta cell-specific expression of mitochondrial GFP (PhAM^flox^Ins1^Cre^). We demonstrate that beta cells transfer mitochondria to islet macrophages *in vivo* and *in vitro*. Diabetogenic stressors did not alter the frequency of mitochondrial transfer and macrophages containing beta cell-derived GFP exhibit increased protein synthesis rates. RNA-seq identified upregulation of activity-regulated cytoskeleton associated protein (*Arc*) in macrophages receiving beta cell-derived mitochondria, while disruption of actin cytoskeleton dynamics prevented mitochondrial transfer. Together, these findings identify mitochondrial transfer as a previously unrecognized mechanism of beta cell-macrophage communication that may contribute to islet homeostasis and immune regulation.

## Introduction

Islet macrophages are key regulators of tissue homeostasis and immune responses in pancreatic islets through antigen presentation and regulation of local inflammatory responses ^1^. Increasing evidence suggests that communication between beta cells and macrophages influences both immune activation and beta cell development and function ^2^. Macrophage function is tightly linked to metabolic status and mitochondria is a key influencer of immune cell behaviour ^3,4^. Pro-inflammatory macrophages upregulate glucose utilisation while shutting down mitochondrial respiration. Alternatively activated macrophages increase glucose uptake upon activation but do so in conjunction with enhanced mitochondrial respiration ^5^.

Type 1 diabetes (T1D) is an autoimmune disease in which autoreactive T cells drive beta cell destruction, while islet macrophages are also recognized to play a role in both beta cell loss and dysfunction and T cell activation in T1D ^1^. In type 2 diabetes (T2D), both insulin resistance and beta cell dysfunction are associated with cytokine production by pro-inflammatory macrophages^6^. In both T1D and T2D, islet inflammation leads to increased expression of HLA class I molecules on the beta cell surface, beta cell endoplasmic reticulum (ER) stress, and mitochondrial ROS production^7–9^, with several recent studies targeting antioxidant responses through reduction of thioredoxin-interacting protein (TXNIP) as an approach to relieve beta cell stress^10–12^. Although T1D and T2D arise through distinct etiologies, in both forms of the disease beta cells undergo chronic cellular stress driven by inflammatory cytokines and characterized by mitochondrial dysfunction, oxidative and ER stress, and impaired bioenergetics^13,14^. These processes not only compromise insulin secretion but may also alter intercellular communication within the islet microenvironment.

It has been recently established that entire mitochondria or their components can be transferred between different cell types, thereby modulating cellular and systemic responses^15–19^. Transfer of mitochondria from bone marrow stromal cells into CD8 T cells enhances antitumor function and metabolic capacity, through increased protein translation rates^19^. In addition, tumor cells hijack mitochondria from immune cells through tunneling nanotube formation, which works as a mechanism of immune evasion by the tumor cells^4,16^. Mitochondria-deficient immune cells show impaired antigen presentation and reduced production of inflammatory mediators, such as cytokines and cytotoxic granules^15^.

Given the evidence for mitochondria transfer as a form of intercellular communication, and the fact that beta cells are highly secretory^20^, we aimed to determine whether mitochondria transfer from beta cells to macrophages occurs in the islet environment. Studies of interactions between beta cells and macrophages are challenging, due to the limited number of islet resident macrophages and highly heterogeneous cell populations. Here, we used a genetic mouse model with beta cell-specific expression of mitochondrial GFP (PhAM^fl^Ins1^cre^), as well as a co-culture model using mitochondrial GFP-expressing insulinoma cells (GFP MIN6) and bone marrow-derived macrophages (BMDMs) to show that mitochondria transfer from beta cells to macrophages occurs physiologically via nanotube formation and is dependent on activity-regulated cytoskeleton-associated protein (*Arc*). Macrophages that receive exogenous mitochondria show increased metabolic activity *in vitro* and *in vivo*. Our data point to a novel mechanism of cross-talk between beta cells and macrophages with potential new targets for regulation of islet macrophage function.

## Results

### Islet macrophages acquire mitochondria from beta cells

To test whether beta cells transfer mitochondria to neighbouring macrophages in pancreatic islets, we first generated photo-activatable mitochondria floxed (PhAM^fl^) mice^21^ crossed with Insulin 1^Cre^ (PhAM^fl^Ins1^Cre^) to obtain mice with beta cell-specific-expression of a mitochondrial GFP insert (mGFP), to track beta cell-derived mitochondrial fate (Fig 1A). Following pancreatic islet isolation and dispersion, we found that >90% of insulin^+^ beta cells were GFP^+^ compared to wild type control (PhAM^fl^, here referred as WT) (Fig 1B). We also induced activation of Cre recombinase in adult mice, using adeno-associated virus serotype 8 (AAV8)-Ins1^Cre^ or empty vector (WT) injection for beta cell recombination^22^ (Supp Fig 1A). We obtained reasonable recombination using PhAM^fl^ mice, with >60% of beta cells expressing GFP, a proportion that was maintained in the presence of cytokine stress (Supp Fig 1B, C). Strikingly, we observed an average of ∼30% of macrophages from these islets to contain beta cell-derived mGFP in both models of PhAM^fl^Ins1^Cre^ recombination (Fig 1C; Supp Fig 1D), suggesting intercellular transfer of mitochondria from beta cells to islet macrophages.

**Figure 1.**
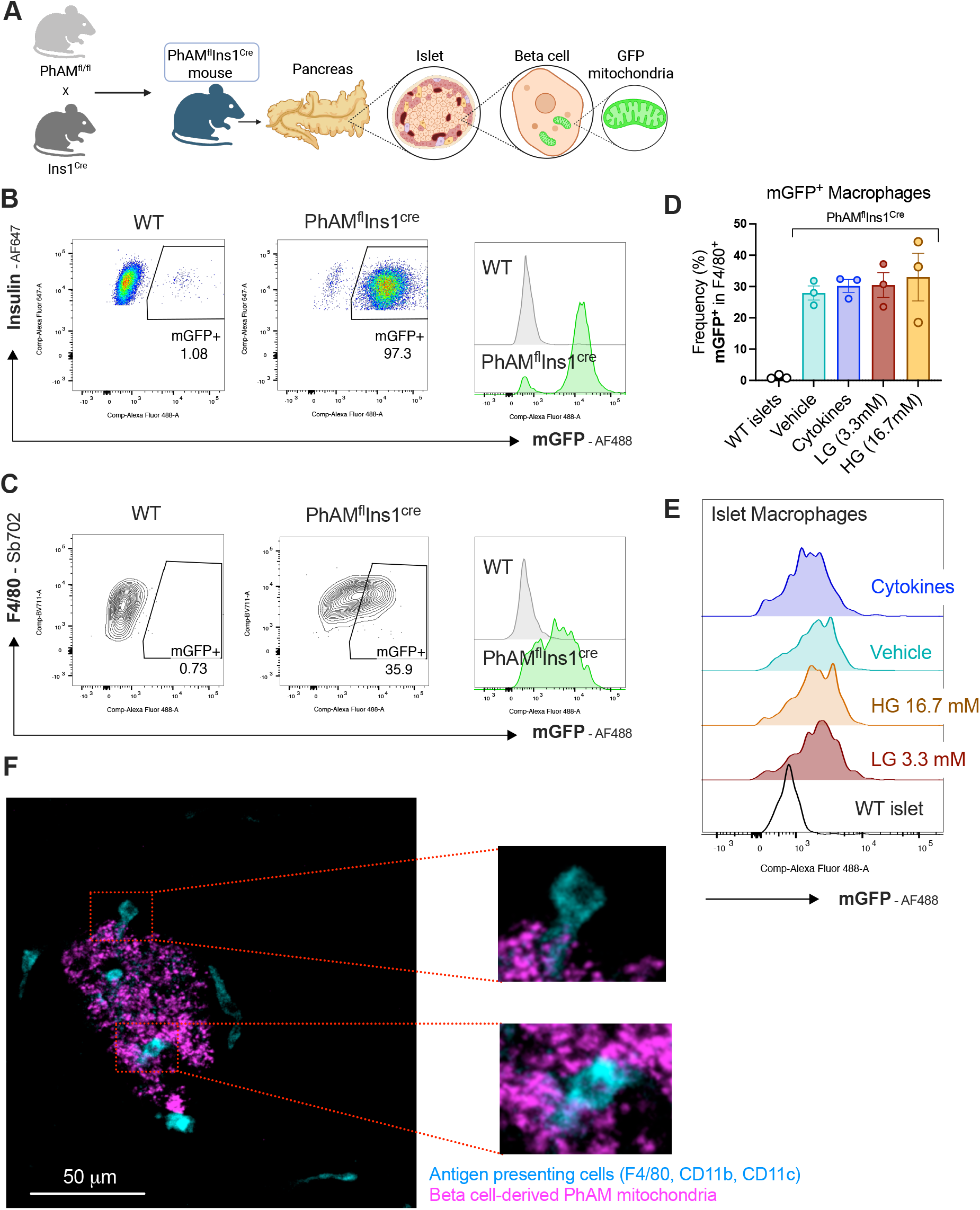
beta cell-derived mitochondrial GFP is present in islet macrophages. Pancreatic islets were collected from Pham^fl^Ins1^Cre^ mice containing a beta cell-specific mitochondrial GFP insert and treated *ex vivo* with either vehicle or pro-inflammatory cytokine cocktail (IL-1β 50 U/mL; TNF-α 1000 U/mL; IFN-γ 1000 U/mL); Low glucose (3.3 mM) or high glucose (16.7 mM). **A)** Schematics depicting breeding strategy for generation of Pham^fl^Ins1^Cre^ mice. **B)** Representative flow plots of mGFP^+^ beta cells (insulin^+^) gated from live cells of whole dispersed islets. **C)** Representative flow plots of mGFP^+^ macrophages (F4/80^+^) gated from live cells of whole dispersed islets. **D)** Percentage of mGFP^+^ macrophages isolated from islets of wild type mice (WT) and Pham^fl^Ins1^Cre^ mice, following treatment with vehicle, pro-inflammatory cytokines, low glucose (LG), or high glucose (HG). **E)** Representative histograms of mGFP^+^ macrophages isolated from islets of wild type mice (WT) and Pham^fl^Ins1^Cre^ mice, following treatment with vehicle, pro-inflammatory cytokines, low glucose (LG), or high glucose (HG). **F)** Representative confocal image of live pancreas showing GFP signal from beta cell-derived mitochondria (pink) and antigen presenting cells (blue) in the islet. Data is shown as mean ± SEM, n ≥ 3 mice.

To investigate how physiologically relevant inflammatory stressors and varying glucose concentrations influence beta cell-macrophage communication, we next exposed intact *ex vivo* WT and PhAM^fl^Ins1^Cre^ islets for 24h to a pro-inflammatory cytokine cocktail, and for 2h to either low (3.3 mM) or high (16.7 mM) glucose. We detected no differences in beta cell mGFP signal when islets were treated *ex vivo* with pro-inflammatory cytokines or varying levels of glucose, with beta-cell derived mGFP-positive macrophage frequency being retained at similar levels (Fig 1D, E; Supp Fig 1E). Next, we sought to determine whether beta cell mGFP is detectable in macrophages in live tissue, without the stress of pancreatic tissue digestion for islet isolation. To this end, we harvested whole pancreata from PhAM^fl^Ins1^Cre^ mice and performed live pancreas imaging. Using focus stacking captures (z-stack), we identified pancreatic islets and captured beta cell mitochondria (mGFP) structures co-localized within the cytoplasmic region of islet antigen-presenting cells (Fig 1F; Supp material video file). Our findings support the hypothesis that mitochondrial transfer is a physiological attribute of islet macrophages that is not dependent on inflammatory cues or nutrient availability.

### Mitochondrial transfer in vitro requires direct cell contact and is not phagocytosis-dependent

To facilitate mechanistic studies of mitochondrial transfer from beta cells to macrophages, we established an *in vitro* co-culture system that recapitulates the phenomenon we observed *in vivo*. While beta cell-derived mGFP could be detected in primary mouse islet macrophages, the low abundance of resident macrophages restricted detailed and functional analyses. In contrast, a reductionist *in vitro* system provides increased cell numbers, enabling comprehensive investigation of the mechanism and functional consequences of mitochondrial transfer. For this purpose, we used the insulinoma cell line MIN6 we engineered to express a mitochondrial matrix-localized roGFP (MitoGFP MIN6)^23,24^ (Fig 2A, Supp Fig 2A, B).

**Figure 2.**
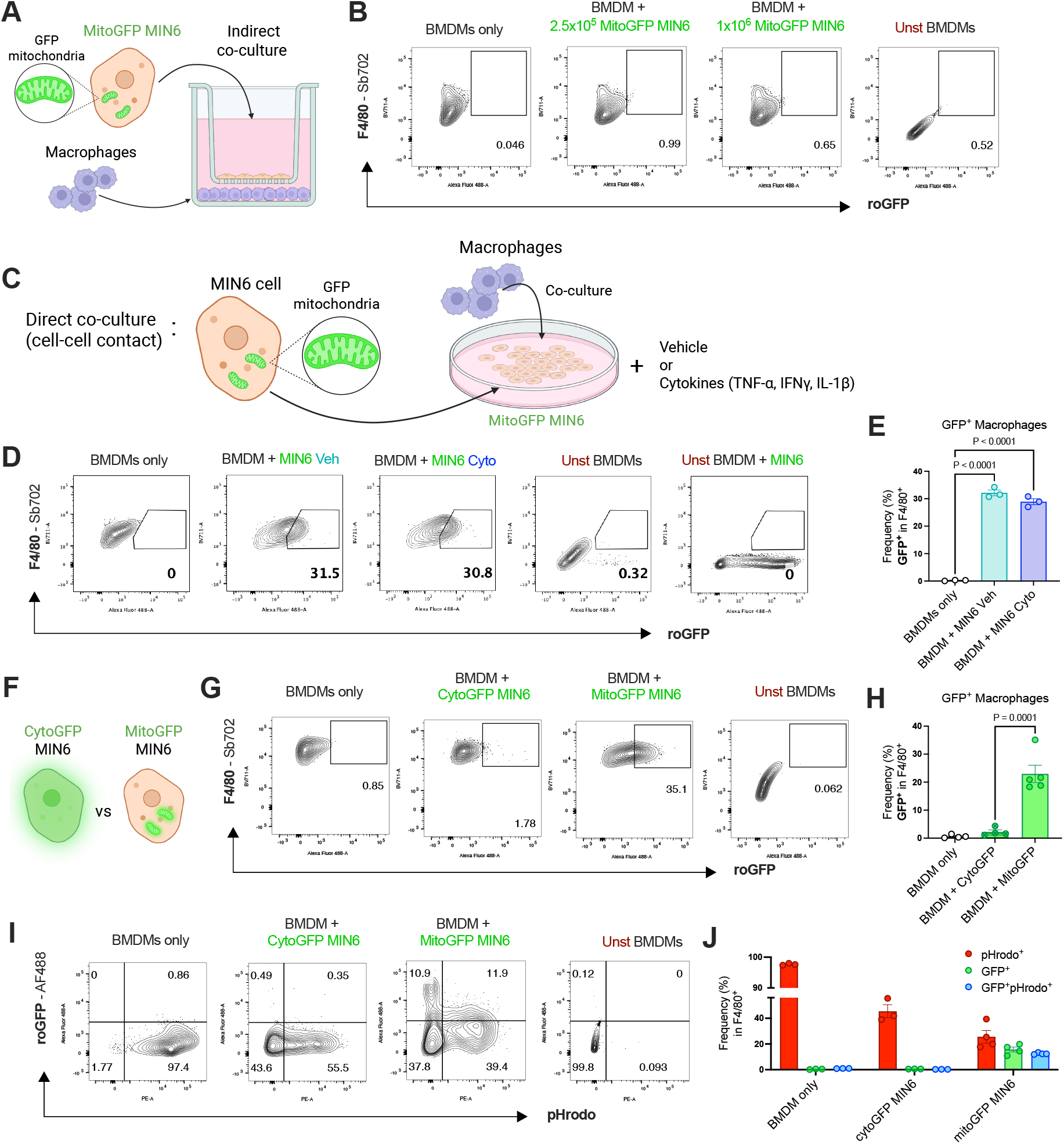
Direct cell contact, but not phagocytosis, is required for mitochondrial transfer. MIN6 beta cells expressing either mitochondrial or cytosolic GFP (MitoGFP; CytoGFP) were co-cultured with BMDMs for assessment of mitochondrial transfer events. **A)** Schematics of indirect co-culture of MitoGFP MIN6 beta cells and BMDMs using Transwell. **B)** Representative flow plots of GFP^+^ macrophages (F4/80^+^) co-cultured with MitoGFP MIN6. Seeding of 2.5 x 10^5^ MIN6 and 1 x 10^6^ MIN6 to 2 x 10^5^ BMDMs. **C)** Schematics of direct cell co-culture of BMDMs and MitoGFP MIN6 beta cells pre-treated with either vehicle or pro-inflammatory cytokines for 18h. Cells were seeded at a 1:5 ratio of BMDM:MIN6. **D)** Representative flow plots c **E)** Frequency of GFP^+^ macrophages (F4/80^+^) alone or co-cultured with MitoGFP MIN6. **F)** Schematics of MIN6 beta cells expressing cytosolic GFP (CytoGFP) or mitochondrial GFP (MitoGFP). **G)** Representative flow plots of GFP^+^ macrophages (F4/80^+^) directly co-cultured with either CytoGFP or MitoGFP MIN6. **H)** Percentage of GFP^+^ macrophages (F4/80^+^) alone or directly co-cultured with either CytoGFP or MitoGFP MIN6. **I)** Representative flow plots of pHrodo^+^ / GFP^+^ macrophages (gated in F4/80^+^) directly co-cultured with either CytoGFP or MitoGFP MIN6. Following co-culture, cells were incubated with pHrodo bioparticles and acquired live by flow cytometry. **J)** Frequency of pHrodo^+^ / GFP^+^ macrophages co-cultured with either CytoGFP or MitoGFP MIN6. Data is shown as mean ± SEM, n ≥ 5 co-cultures. P values < 0.05 are considered significant.

To determine whether mitochondria are released from MIN6 beta cells and taken up by neighbouring macrophages, we performed indirect co-culture, seeding bone marrow-derived macrophages (BMDMs) into a cell culture plate and adding MitoGFP MIN6 cells into a Transwell cell culture system to enable media content exchange (Fig 2A). Using this indirect co-culture model, we did not detect any beta cell-derived GFP in BMDMs (Fig 2B), even after increasing the ratio of MIN6 cells to BMDMs (Fig 2B). We then asked whether cell contact was necessary for mitochondrial transfer. Direct co-culture of MitoGFP MIN6 + BMDMs resulted in detection of GFP^+^ in an average of ∼30% of BMDMs (Fig 2C, D), much like the ex-vivo analysis of islet-resident macrophages (Fig 1D). When MitoGFP MIN6 cells were pre-treated for 18 h with vehicle or pro-inflammatory cytokine cocktail (TNF-α, IL-1β, IFN-γ), the frequency of GFP^+^ BMDMs was not altered upon contact (Fig 2D, E). We also rested MitoGFP MIN6 cells in low glucose (3.3 mM) for 1 h before incubating cells in either low or high (16.7 mM) glucose for 2 h. We did not detect any changes in the frequency of GFP^+^ macrophages following high glucose culture, with a frequency of ∼30% of BMDMs becoming GFP^+^ following co-culture (Supp Fig 2C, D). These data show that BMDMs acquire beta cell-derived mitochondrial GFP independent of inflammatory stimuli, or glucose deprivation and glucose level spikes.

Next, we sought to understand whether the presence of GFP in BMDMs after co-culture was a specific mechanism related to direct transfer of mitochondria from beta cells or derived from beta cell granules or macrophage sampling of cytosol-containing vesicles^25,26^. We used MIN6 cells we engineered to express cytosolic roGFP (CytoGFP) or mitochondrial roGFP (MitoGFP)^23,24^ and co-cultured these cells with BMDMs (Fig 2F). We found that only MitoGFP was present in BMDMs following co-culture (Fig 2G, H), indicating that GFP^+^ macrophages are a result of mitochondrial transfer and not due to macrophage acquisition of other cellular components. We also aimed to determine whether the presence of GFP in BMDMs could occur via particle engulfment during phagocytosis. For this, we incubated Cyto-or MitoGFP MIN6 + BMDMs with pHrodo *E. coli* bioparticles, which fluoresce upon acidification in the phagolysosome. Consistent with our previous results, we did not observe GFP^+^ macrophages when co-cultured with CytoGFP MIN6 (Fig 2I, J). We found distinct macrophage populations performing separate roles: GFP^+^ macrophages that received exogenous mitochondria from beta cells; pHrodo^+^ phagocytic macrophages; and GFP^+^pHrodo^+^ that are both phagocytic and engaged in mitochondrial transfer from beta cells (Fig 2I, J; Supp Fig 2E, F). These distinct macrophage population were present even 72h after co-culture (Fig 2I, J; Supp Fig 2E, F). Together, these findings highlight mitochondrial transfer from beta cells to macrophages as a unique biological process that is maintained even in long culture times.

### Acquisition of exogenous mitochondria alters macrophage metabolic capacity

To evaluate potential functional changes resulting from acquisition of exogenous beta cell-derived mitochondria, we assessed single cell energetic metabolism by measuring ATP-dependent protein translation via SCENITH^27,28^. We used both primary islets isolated from PhAM^fl^Ins1^Cre^ mice, as well as GFP MIN6 + BMDMs co-cultured cells (Fig 3A). In both models, macrophages showed sensitivity to both glycolysis inhibitor 2-deoxy-D-glucose (2-DG) and ATP synthase inhibitor oligomycin (Oligo) (Fig 3B, C). Interestingly, GFP^+^ macrophages showed overall higher protein translation rates compared to GFP^-^macrophages even under the effect of metabolic inhibitors (Fig 3B, C). We next evaluated mitochondrial membrane polarization states in GFP^+^ and GFP^-^macrophages. Mitochondrial membrane potential measured by both tetramethylrhodamine methyl ester (TMRM) and MitoTracker Red CMXRos (MT Red CMXRos) revealed that GFP^+^ macrophages exhibited higher mitochondrial membrane potential compared to GFP^-^macrophages (Fig 3D, E). Additionally, GFP^+^ macrophages displayed increased expression of co-stimulatory molecule CD86 under basal conditions and increased expression of TNFα following pro-inflammatory cytokine treatment (Fig 3F, G). Collectively, these results suggest that acquisition of exogenous beta cell-derived mitochondria may support macrophage metabolic capacity, pro-inflammatory, and antigen presenting function.

**Figure 3.**
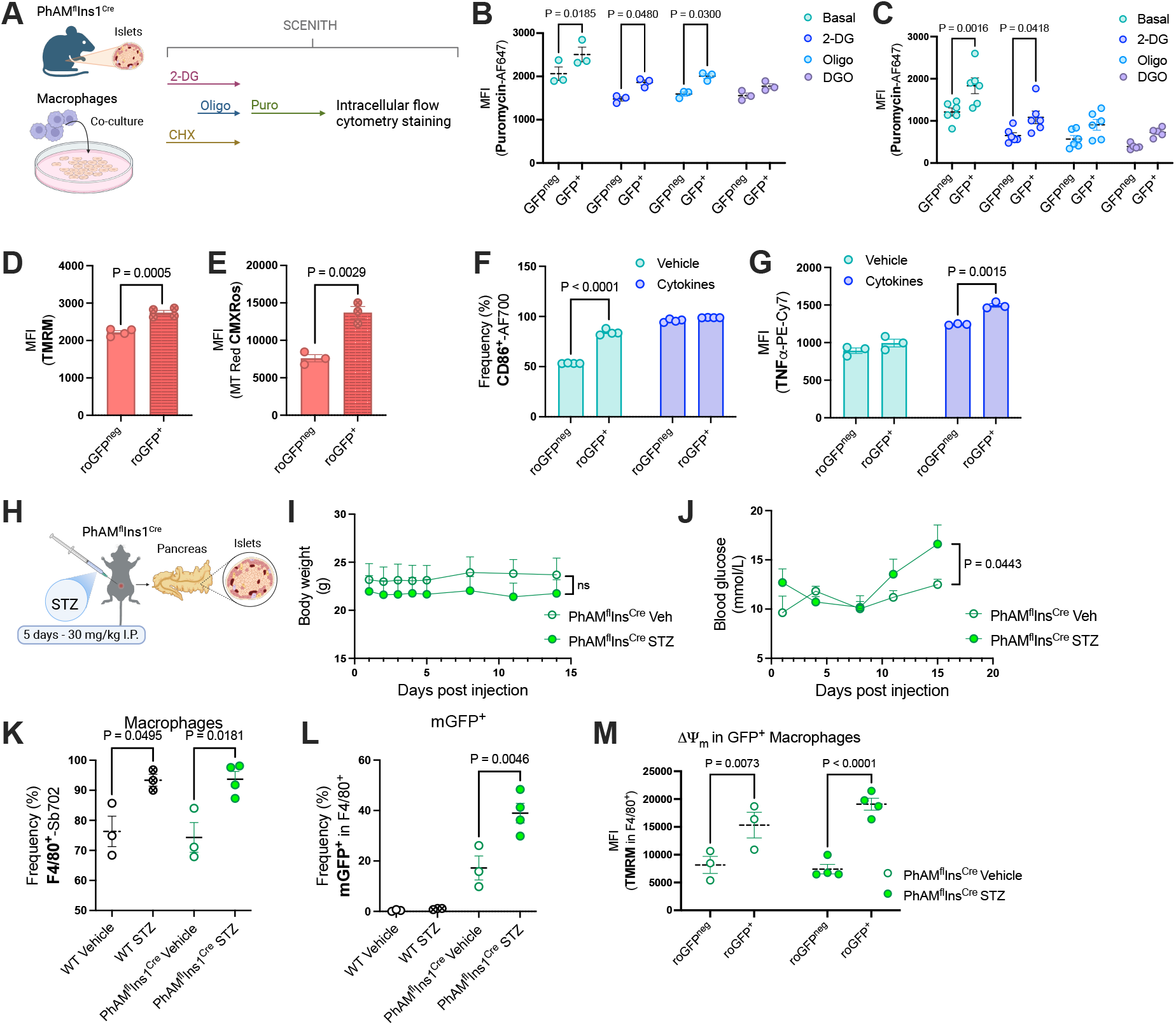
Uptake of beta cell-derived mitochondria modulates macrophage metabolism. Pancreatic islets isolated from Pham^fl^Ins1^Cre^ mice as well as MitoGFP MIN6 + BMDMs co-cultures were used for analyses. **A)** Schematics of SCENITH assay from dispersed islets of Pham^fl^Ins1^Cre^ mice and MitoGFP MIN6 + BMDMs co-cultures. **B)** Mean fluorescence intensity (MFI) of puromycin in GFP^+^ and GFP^-^ macrophages from MitoGFP MIN6 + BMDMs co-cultures at basal levels, and following treatment with 2-deoxyglucose (2-DG), oligomycin (Oligo), or 2-G + Oligo (DGO). **C)** MFI of puromycin in GFP^+^ and GFP^-^macrophages from primary islets of Pham^fl^Ins1^Cre^ mice at basal levels, and following treatment with 2-DG, Oligo, or DGO. **D)** MFI of TMRM in GFP^+^ and GFP^-^ macrophages, indicating mitochondrial membrane potential. **E)** MFI of MitoTracker (MT) Red CMXRos in GFP^+^ and GFP^-^ macrophages, indicating mitochondrial membrane potential. **F)** Frequency of CD86 expression in GFP^+^ and GFP^-^ macrophages following co-culture with MitoGFP MIN6 pre-treated with either vehicle or pro-inflammatory cytokines. **G)** Frequency of TNF-α expression in GFP^+^ and GFP^-^ macrophages following co-culture with MitoGFP MIN6 pre-treated with either vehicle or pro-inflammatory cytokines. **H)** Schematic representation of multiple low-dose streptozotocin (STZ)-induced beta cell dysfunction. WT and Pham^fl^Ins1^Cre^ mice were treated with 30 mg/kg of STZ for 5 consecutive days and islets were harvested 10 days after the last STZ injection. **I)** Body weight of Pham^fl^Ins1^Cre^ mice treated with vehicle or STZ. **J)** Blood glucose of Pham^fl^Ins1^Cre^ mice treated with vehicle or STZ. **K)** Percentage of macrophages (F4/80^+^) in WT and Pham^fl^Ins1^Cre^ mice treated with vehicle or STZ. **L)** Percentage of mGFP^+^ macrophages (F4/80^+^) in WT and Pham^fl^Ins1^Cre^ mice treated with vehicle or STZ. **M)** MFI of TMRM in GFP^+^ and GFP^-^ macrophages of Pham^fl^Ins1^Cre^ mice treated with vehicle or STZ. Data is shown as mean ± SEM, n ≥ 5 co-cultures; n ≥ 3 mice per experiment. P values < 0.05 are considered significant.

Macrophage frequency increases in the islet following beta cell stress and death, accompanied by metabolic adaptations linked to oxidative phosphorylation and lysosomal activity^29^. We submitted healthy adult PhAM^fl^Ins1^Cre^ and WT control mice to 5 consecutive daily intraperitoneal (i.p.) injections of low dose streptozotocin (STZ) to induce beta cell death *in vivo* (Fig 3H). Islets were harvested 10 days after the last STZ injection. STZ-treated mice displayed no changes in body weight (Fig 3I; Supp Fig 3A) but exhibited mild hyperglycemia10 days after the last STZ injection (Fig 3J; Supp Fig 3B). Islets isolated from STZ-treated mice showed higher macrophage frequency (Fig. 3K). Notably, STZ-treated PhAM^fl^Ins1^Cre^ mice had an increased frequency of mGFP^+^ macrophages compared to vehicle-treated mice (Fig 3L). Similar to our *in vitro* findings, mGFP^+^ islet macrophages had higher mitochondrial membrane potential in both vehicle and STZ-treated conditions. Notably, using an *in vivo* approach of beta cell stress and death led to accumulation of mGFP signal in islet macrophages that was not observed in *in vitro* models of inflammation. Together, these data identify beta cell-derived mitochondrial transfer as a mechanism that improves the metabolic fitness of recipient macrophages, supporting a functional role for mitochondrial exchange within the islet microenvironment.

### Mitochondrial transfer requires actin cytoskeleton remodelling

Tunnelling nanotube formation is one of the mechanisms by which cells exchange mitochondria *in vitro* and *in vivo*^15,16,19,30^. We observed that co-cultured cells form mitochondria-rich tubular structures bridging beta cells and macrophages in *in vitro* co-cultures (Fig 4A). To understand the mechanism that drives nanotube structure formation and mitochondrial acquisition by macrophages, we sorted GFP^+^ and GFP^-^macrophages after co-culture with either vehicle-or cytokine-treated MitoGFP MIN6, followed by bulk mRNA sequencing (Fig 4B). Gene expression profiling revealed principal component (PC) 1 and PC2 with 93% and 3% variance, respectively (Supp Fig 4A). The biggest variance observed was between vehicle-and cytokine-treated groups, followed by GFP^+^ and GFP^-^groups, specified by the presence of beta cell-derived mitochondrial GFP (Supp fig 4A, B). According to DE-Seq2 analysis, 837 genes were upregulated and 441 downregulated in GFP^+^ vs. GFP^-^ vehicle-treated macrophages (thresholds of log_2_ fold change >0.25 per comparison) out of 1278 differentially expressed genes. In cytokine-treated GFP^+^ vs. GFP^-^ macrophages, 125 genes were upregulated and 255 genes were downregulated out of 380 differentially expressed genes. Transcriptomic KEGG enrichment analysis of upregulated genes in GFP^+^ macrophages indicated pathways related to MAPK signalling pathway and regulation of actin cytoskeleton in vehicle-treated conditions (Supp Fig 4C). In cytokine-treated conditions, amongst the most upregulated pathways in GFP^+^ macrophages, were cytokine-cytokine receptor interactions and MAPK signalling pathway (Supp Fig 4D). Several complex cellular programs are regulated by MAPK families, including cell development, differentiation, and proliferation^31^. Cell-cell contact has been shown to reduce protein expression of cJun (a transcription factor target of MAP kinase JNK) but higher phosphorylation of p38 and ERK1/2^32,33^. Consistent with pathway enrichment highlighting regulation of actin cytoskeleton, the activity-regulated cytoskeleton-associated protein (*Arc*) was found to be one of the highest commonly upregulated gene in GFP^+^ macrophages from both vehicle-and cytokine-treated co-cultures (Fig 4C).

**Figure 4.**
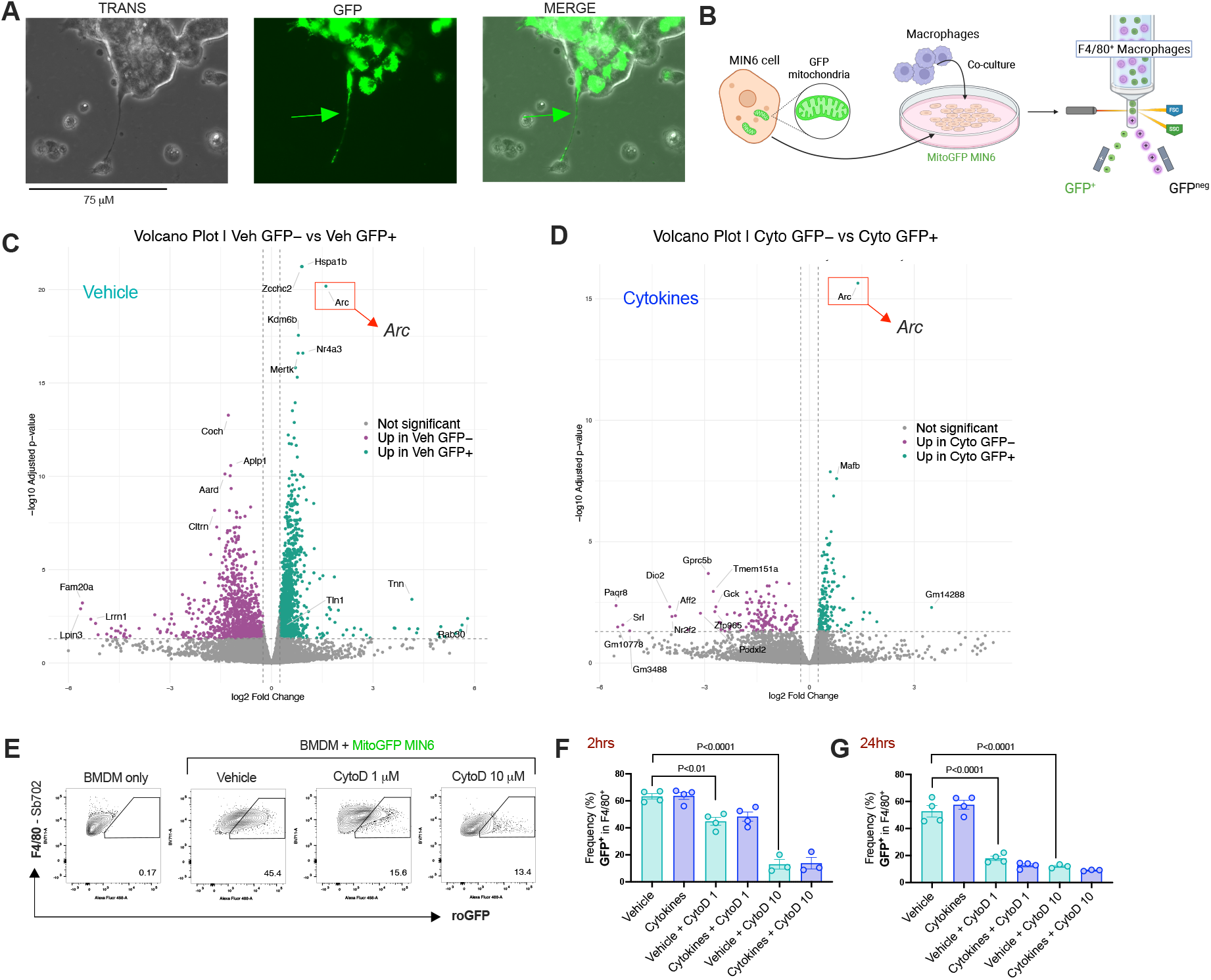
Arc-dependent actin cytoskeleton remodelling mediates mitochondrial transfer. Following co-culture with MitoGFP MIN6 beta cells, GFP^+^ and GFP^-^BMDMs (F4/80^+^) were sorted for RNA isolation and sequencing. **A)** Representative fluorescent microscopy image of MitoGFP MIN6 beta cells co-cultured with BMDMs. Arrow indicates mitochondria-rich nanotube. **B)** Schematics of MitoGFP MIN6 + BMDMs co-culture followed by cell sorting of GFP^+^ and GFP^-^ BMDMs for transcriptomic analysis. **C)** Volcano plot highlighting upregulated genes in GFP^+^ macrophages co-cultured under basal (vehicle) condition. **D)** Volcano plot highlighting upregulated genes in GFP^+^ macrophages co-cultured under beta cell pro-inflammatory cytokine stress. **E)** Representative flow plots of GFP^+^ macrophages (F4/80^+^) after treatment with either vehicle, 1 or 10 μM of cytochalasin D (CytoD). **F)** Frequency of GFP^+^ macrophages (F4/80^+^) from 2h of co-cultures pre-treated with vehicle or pro-inflammatory cytokines and followed by 1 or 10 μM of CytoD. **G)** Frequency of GFP^+^ macrophages (F4/80^+^) from 24h of co-cultures pre-treated with vehicle or pro-inflammatory cytokines and followed by 1 or 10 μM of CytoD. Data is shown as mean ± SEM, n ≥ 4 co-cultures. P values < 0.05 are considered significant.

Tunnelling nanotube formation can be disrupted by the mycotoxin cytochalasin B and inhibit intercellular organelle transfer^34^. Cytochalasin binds to actin filaments and prevents actin cytoskeleton polymerization. We used cytochalasin D to avoid effects related to inhibition of glucose transporters observed with cytochalasin B^35^. We treated co-cultured GFP MIN6 + BMDMs with 1 or 10 µM cytochalasin D (CytoD) and assessed mitochondrial transfer rates. CytoD effectively prevented mitochondrial transfer to macrophages at both concentrations (Fig 4F, G). Inhibition of mitochondrial transfer by CytoD was observed after 2 h of treatment and was more pronounced at 24 h, almost completely blocking mitochondrial transfer (Fig 4G, H). These findings indicate that mitochondrial transfer between beta cells and macrophages occurs through actin cytoskeleton remodelling.

## Discussion

Here, we establish mitochondrial transfer from beta cells to macrophages as a mechanism of intercellular communication that modulates islet macrophage metabolism and inflammatory function, highlighting intercellular organelle exchange as a previously unappreciated component of immune-endocrine crosstalk in pancreatic islets. The recent discovery of intercellular and inter-organ mitochondria transport has stimulated research into the biology of mitochondrial transfer and its therapeutic potential. Restoration of respiration and ATP production can be achieved by transplanting healthy mitochondria into tissue affected by acute injury or cancer^36–39^. In the pancreas, macrophages play a pivotal role in tissue homeostasis but are also drivers of chronic inflammation and beta cell death through release of pro-inflammatory cytokines, which can affect insulin secretion and sensitivity^40–43^. A tightly regulated balance between beta cell and macrophage function is necessary for proper systemic glycemic control.

Our data show that transfer of mitochondria from beta cells to neighbouring macrophages requires direct cell-cell contact, since we did not detect presence of fluorescently labelled mitochondria in recipient cells using a Transwell system. This finding fits with other reports of mitochondrial transfer between cells, including between tumor and immune cells, requiring cell-cell contact^16^. Interestingly, the rates of mitochondrial transfer did not change in the presence of beta cell stimuli or stresses seen in the pancreatic islet microenvironment in diabetes, including high glucose and cytokines. Pancreatic beta cells are expert glucose sensors, allowing dynamic glycemic control through glucokinase activity^44^. When treated with low or high glucose, mitochondrial transfer rates remained unchanged in both primary islets and in a co-culture model using a beta cell line with mitochondrial GFP expression and macrophages. The same pattern was observed in islet and cell cultures treated with pro-inflammatory cytokines. We found that pro-inflammatory cytokine treatment increased the expression of TNF-α and the co-stimulatory molecule CD86 in macrophages but GFP^+^ macrophages showed even higher expression, a finding observed even in vehicle-treated cells. These findings suggest that mitochondrial transfer could have a role in the interaction between islet macrophages and T cell activation. Similar findings have highlighted a role for exogenous mitochondria uptake in increased pro-inflammatory function, in which CD8 T cells that received mitochondria from bone marrow stromal cells showed improved antitumor function and lower expression of exhaustion markers^19^. Overall, our findings suggest that mitochondrial transfer is a physiological and ubiquitous process that modulates macrophage inflammatory markers and metabolic activity.

Macrophage activation and function rely on their core metabolic programming^5,45,46^. Through SCENITH analysis of protein translation rates, we found that both primary islet macrophages and BMDMs co-cultured with MIN6 beta cells with mitochondrial GFP expression showed sensitivity to glycolysis inhibition with 2-DG and ATP synthase inhibition with oligomycin. GFP^+^ macrophages displayed overall higher protein translation rates (as indicated by increased puromycin incorporation) and higher mitochondrial membrane potential than GFP^-^ macrophages. In other studies, recipient cells have similarly been shown to have increased metabolic capacity upon acquisition of exogenous mitochondria^16,19^, proposed to be an adaptive mechanism in situations of high metabolic demand or low nutrient availability.

Another interesting observation is during beta cell dysfunction and diabetes development. Streptozotocin (STZ) treatment increases antigen presentation via MHC class II and upregulation of CD86 expression in macrophages^47,48^. In addition, islet macrophages from STZ-treated mice display higher frequency of macrophages containing beta cell-derived mitochondria and show higher expression of CD86. In primary islets isolated from STZ-treated mice, the frequency of macrophages that receive beta cell mitochondria is higher compared to vehicle-treated mice; whereas induction of beta cell stress *in vitro* did not alter mitochondrial transfer event rates. This suggests that mitochondrial transfer between beta cells and macrophages is not only linked to beta cell inflammatory stress, as *in vivo* induction of beta cell dysfunction through STZ led to higher rates of mitochondrial transfer. STZ-induced beta cell death and dysfunction induce macrophage expression of insulin-like growth factor 1 and genes involved in oxidative phosphorylation (OXPHOS)^29^. High rates of mitochondrial transfer coupled with enhanced OXPHOS in STZ-treated islets align with higher protein synthesis rates observed in macrophages that acquire beta cell-derived mitochondria, even after inhibition of glycolysis and ATP synthase by 2-DG and oligomycin.

Several mechanisms have been implicated in intercellular mitochondrial transfer, including tunnelling nanotubes^16,17,19,34,49^, gap junctions^37,50^, endocytosis^51^, extracellular vesicles^52^, and the uptake of free extracellular mitochondria by recipient cells^53–55^. These complementary pathways enable mitochondrial exchange across a variety of physiological and pathological settings and may be preferentially engaged depending on the donor and recipient cell types. Here, we show that *Arc* (Activity-Regulated Cytoskeleton-associated protein) is upregulated in GFP^+^ macrophages, which acquired mitochondria from beta cells. *Arc* is a neuronal plasticity protein that is also involved in dendritic cell migration^56^. We identified tubular elongations in co-cultures of beta cells and macrophages, which indicates that *Arc*-mediated cytoskeleton remodelling is required for tubular structure formation^57^. In line with this, nanotube-mediated mitochondrial transfer can be disrupted by cytochalasin B^17^ and cytochalasin D, which shows that polymerization of actin filaments is required for nanotube-mediated mitochondrial transfer between beta cells and macrophages.

Taken together, our findings demonstrate that pancreatic beta cells transfer mitochondria to macrophages through a nanotube-dependent mechanism that enhances mitochondrial membrane potential, metabolic activity, and cytokine production in recipient cells, identifying mitochondrial transfer as a functional mechanism of beta cell-macrophage communication within the islet microenvironment.

### Limitations

Mitochondrial transfer events from beta cells to macrophages are rare in primary islets, due to the scarcity of islet-resident macrophages, which limited the number of recipient cells available for downstream analyses. This prompted the use of complementary *in vitro* models to investigate the functional consequences of mitochondrial transfer. In addition, although our findings focused on the transfer of mitochondria from beta cells to macrophages, we do not discard the hypothesis that mitochondrial exchange can be a bidirectional process. Whether macrophages also donate mitochondria to beta cells, and the functional significance related to this, remains to be determined. Finally, our study was performed under defined experimental conditions, and future studies using additional models of diabetes, including autoimmune models such as the NOD mouse and metabolic disease models with high fat diet-induced diabetes, will be important to establish the prevalence, regulation, and physiological relevance of beta cell-macrophage mitochondrial transfer across distinct contexts of diabetes.

## Materials and Methods

### Mice

C57BL/6 mice purchased from the Jackson Laboratory were housed at the BC Children’s Hospital Research Institute Animal Facility with *ad libitum* access to water and chow diet (6% fat, Teklad 2918), under experimental protocol #A24-0113, according to standing guidelines. Female and male mice aged 10-20 weeks were used. Beta cell-specific expression of mitochondrial Dendra2, a photoconvertible GFP protein (mitoGFP), were generated by crossing PhAM floxed (PhAM^fl^) mice (Jackson B6;129S-*Gt(ROSA)26Sor^tm1(CAG-COX8A/Dendra2)Dcc^*/J) with Insulin1 Cre (Ins1^Cre^) mice (Jackson B6(Cg)-*Ins1^tm1.1(cre)Thor^*/J) and by self-complementary adeno-associated virus 8 (scAAV8)-Ins1^Cre^ hepatopancreatic intraductal injection (doi: 10.21769/BioProtoc.5657).

### AAV8-Ins1^Cre^ delivery

AAV8-Ins1^Cre^ and AAV8-Ins1 empty (control) were used at 10^11^ viral genomes (vg) per mouse from stock of 10^13^ vg/mL (Vector Biolabs *pAAV210119-1011afm and VB230512-1203mkx)*. Fresh sterile-filtered AAV8 buffer (0.1% fast green dye, 0.001% pluronic F-68 non-ionic surfactant, in PBS) was used for preparation of AAV8 working solution. AAV8 solution was directly injected into the hepatopancreatic duct of PhAM^fl^ mice at 6-8 weeks of age^58,59^.

### Mouse pancreatic islets

Collagenase solution was freshly prepared by adding 1000 U/mL of collagenase XI (#C7657; Sigma) in Hank’s Balanced Salt Solution (HBSS). Collagenase solution was injected via the common bile duct (1-2 mL). Pancreas was collected and further digested in 50 mL Falcon tube containing collagenase solution for 15 min at 37 °C with shaking. Islet media (1640 RPMI glucose +, L-glutamine + [#10-040-CV; Corning], supplemented with 10% foetal bovine serum [FBS, #12483020; Thermo Fisher Scientific], 1% penicillin/streptomycin [P/S, #SV30010; Thermo Fisher Scientific], and 1% GlutaMAX [#35050061; Gibco]) was used to stop digestion. Islets were filtered through a 70 µm cell strainer and hand-picked for overnight incubation in islet media for recovery and further experimentation^60^. For *ex vivo* treatment, islets were cultured for 18 hours with vehicle (cell culture-grade water) or pro-inflammatory cytokines (interleukin [IL]-1β 50 U/mL, Peprotech #211-11B; TNF-α 1000 U/mL, Peprotech #315-01A; Interferon-γ [IFN-γ] 1000 U/mL, Peprotech #315-05) or pre-incubated for 1 hour with low glucose (LG) media (3.3 mM glucose) followed by incubation for 2 hours in either low glucose (LG 3.3 mM) or high glucose (HG 16.7 mM) media. For islets dispersion into cell suspensions, islets were transferred into FACS tubes (Falcon^®^ round-bottom polystyrene tubes, 5mL, #CA60819-820; VWR) containing 500 mL of PBS + 2.5 mM EDTA and centrifuged at 300 x g for 1 min at room temperature (RT). Supernatant was gently removed and 300 µL of TrypLE Express (#12604021; Gibco) was added to each tube and incubated at 37 °C for dispersion. Islets were vigorously resuspended by pipetting up and down every 3 min until cell suspension was homogeneous and no visible islets were present. FACS buffer (PBS + 2% FBS) was added at 3x the volume of TrypLE to stop the digestion. Cells were then centrifuged at 400 x g for 5 min at 4 °C and supernatant was removed for flow cytometry staining, described below.

### Streptozotocin-induced beta cell damage

Streptozotocin (STZ, #S0130; Sigma) was calculated for each mouse based on body weight, as previously described^28^. STZ was prepared fresh in 0.2 M sodium acetate solution (pH 4.5) and used within 5 min after resuspension. Mice received daily intraperitoneal injections of STZ (30 mg/kg of body weight) for 5 consecutive days. All STZ aliquots were neutralized in 5% KOH solution for 24 hours before being discarded. Mice were monitored for changes in activity, body weight, and blood glucose levels via tail vein poke. Pancreatic islets were harvested 10 days after the last STZ injection for further analysis.

### L929-conditioned media

L929 cells were used as a source of macrophage colony stimulating factor (M-CSF) for bone marrow-derived macrophage (BMDM) differentiation. L929 cells were cultured in T182 flasks containing 50 mL RPMI supplemented with 10% FBS, 1% P/S for 10 days. M-CSF-rich media was collected and centrifuged for pelleting of debris. Supernatant was then sterile filtered (0.2 µm) and titrated for optimal BMDM differentiation, assessed by flow cytometry (expression of F4/80 and CD11b), described below.

### Bone marrow-derived macrophages

Femur and tibia were harvested from C57BL/6 mice, and the bone marrow was flushed from the bone cavity after removal of the epiphyses. Red blood cells were lysed with ammonium chloride buffer (150 mM ammonium chloride, 10 mM potassium bicarbonate, 0.1 mM EDTA, pH 7.3-7.4) for 3 min. The remaining cells were then plated in 1640 RPMI media containing 10% FBS, 1% P/S, 1% HEPES (#SH3023701; Thermo Fisher Scientific), and 20% L929-conditioned media, for 7 days. Media was changed on days 2 and 4. On day 7, fully differentiated macrophages were lifted from the plate using PBS + 2.5 mM EDTA and gently scraped with cell scrapers.

### MIN6 cells

Mouse insulinoma MIN6 cells were cultured in Addex Bio Advanced DMEM medium (#C0003-04; Cedarlane) supplemented with 15% FBS, 1% P/S, and 0.05 mM 2-mercaptoethanol (#21985023; Invitrogen). Expression of cytosolic or mitochondrial GFP (cytoGFP, mitoGFP) in MIN6 cell was achieved by viral transduction, described below.

### Virus generation and transduction

Cytosolic GFP (CytoGFP) and mitochondrial GFP (MitoGFP) construct sequences (Addgene plasmid #49435 and #49437) were cloned into the MSCV backbone (MIGR1; Addgene plasmid #27490) following digestion with Bglll and Sall to remove the IRES-GFP cassette, and ecotropic retrovirus was generated in 293T cells, as previously described (DOI: 10.1038/nature08097). MIN6 cells were transduced with supernatant containing GFP-expressing MSCV retrovirus in the presence of polybrene (8 µg/mL) for 12 hours.

### Co-culture model

BMDMs and GFP MIN6 were co-cultured at a ratio of 1:5 (BMDM:GFP MIN6). For indirect co-culture, BMDMs were seeded and GFP MIN6 cells were added into transwell cell culture inserts (1 µm pore size, #10769-228; VWR). For direct co-culture, BMDMs and GFP MIN6 were seeded into cell culture wells allowing cell-cell contact. GFP MIN6 cells were pre-treated for 18 hours with either vehicle (cell culture-grade water) or pro-inflammatory cytokines cocktail (interleukin [IL]-1β 50 U/mL, Peprotech #211-11B; TNF-α 1000 U/mL, Peprotech #315-01A; Interferon-γ [IFN-γ] 1000 U/mL, Peprotech #315-05). BMDMs were then added at 1:5 ratio for 2 hours. Cells were then harvested for further analysis. GFP MIN6 cells were cultured in low glucose (LG) media (3.3 mM glucose) for 1 hour and then incubated with either LG (3.3 mM glucose) or high glucose (HG, 16.7 mM glucose) with addition of BMDMs (1:5 ratio) for 2 hours. Cells were then collected for further analysis. For mitochondrial transfer inhibition experiments, co-cultured cells were with either vehicle (DMSO), 1 µM, or 10 µM Cytochalasin D (#C8273, Sigma) for either 2 hours or 24 hours.

### Flow cytometry

Co-cultured GFP MIN6 + BMDMs and primary islet cell suspensions were collected in ice cold FACS buffer and the leftover cell cultured media was washed out by centrifugation (400 x g for 5 min at 4 °C). For cell surface markers and viability dye, cells were resuspended in the appropriate antibody cocktail mix (Table 1) and incubated at 4 °C for 20 min in the dark. Cells were washed again with FACS buffer and resuspended in FACS buffer for acquisition in flow cytometer or further intracellular staining was performed. For intracellular staining, cells were resuspended in eBioscience™ Foxp3/Transcription Factor Fixation/Permeabilization Concentrate and Diluent (#00-5521-00, Invitrogen), following manufacturer’s instructions and incubated for 20 min at 4 °C in the dark. Cells were washed using Perm buffer 1x (#00-8333-56, Invitrogen) and centrifugating for 5 min at 400 x g. Intracellular antibody cocktail mix was prepared in Perm buffer 1x (Table 1). Cells were incubated in intracellular antibody cocktail mix at 4 °C for 40 min in the dark. Cells were then washed once again in perm buffer and resuspended in FACS buffer for acquisition in flow cytometer. For live cell acquisition with MitoProbe TMRM (#M20036; Thermo Fisher Scientific), or pHrodo Red E. coli Bioparticles (#P35361; Thermo Fisher Scientific), the dyes/bioparticles were prepared in warm FACS buffer and incubated for 30 min (TMRM) or 15 min (pHrodo) at 37 °C, following the manufacturer’s instructions. Cells were washed and resuspended in FACS buffer for immediate acquisition. Details for antibodies and probes dilutions are listed in Table 1. Acquisition was performed using a BD FACS Symphony A1. Data were analysed using FlowJo Software license (BD).

**Table 1.**
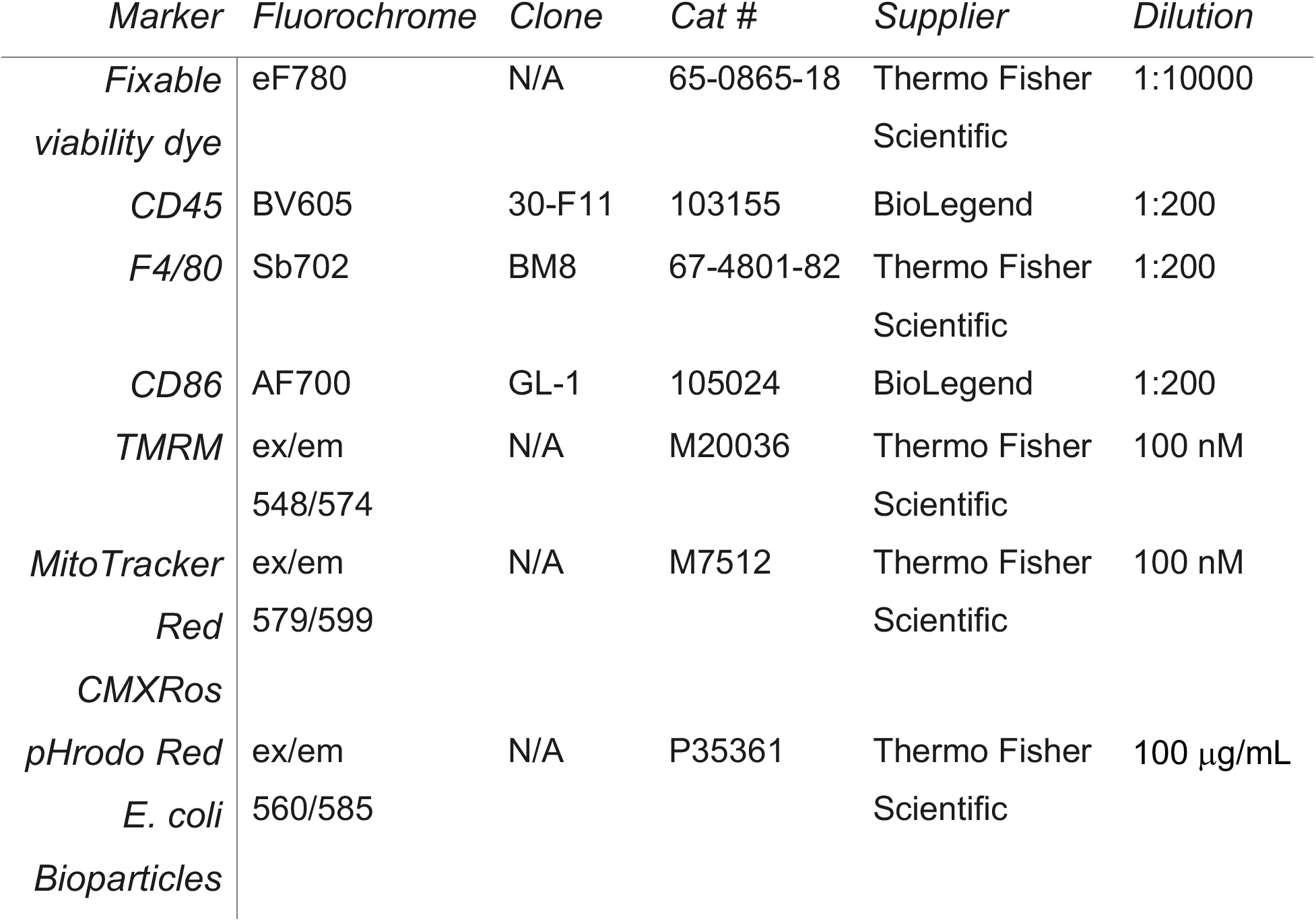
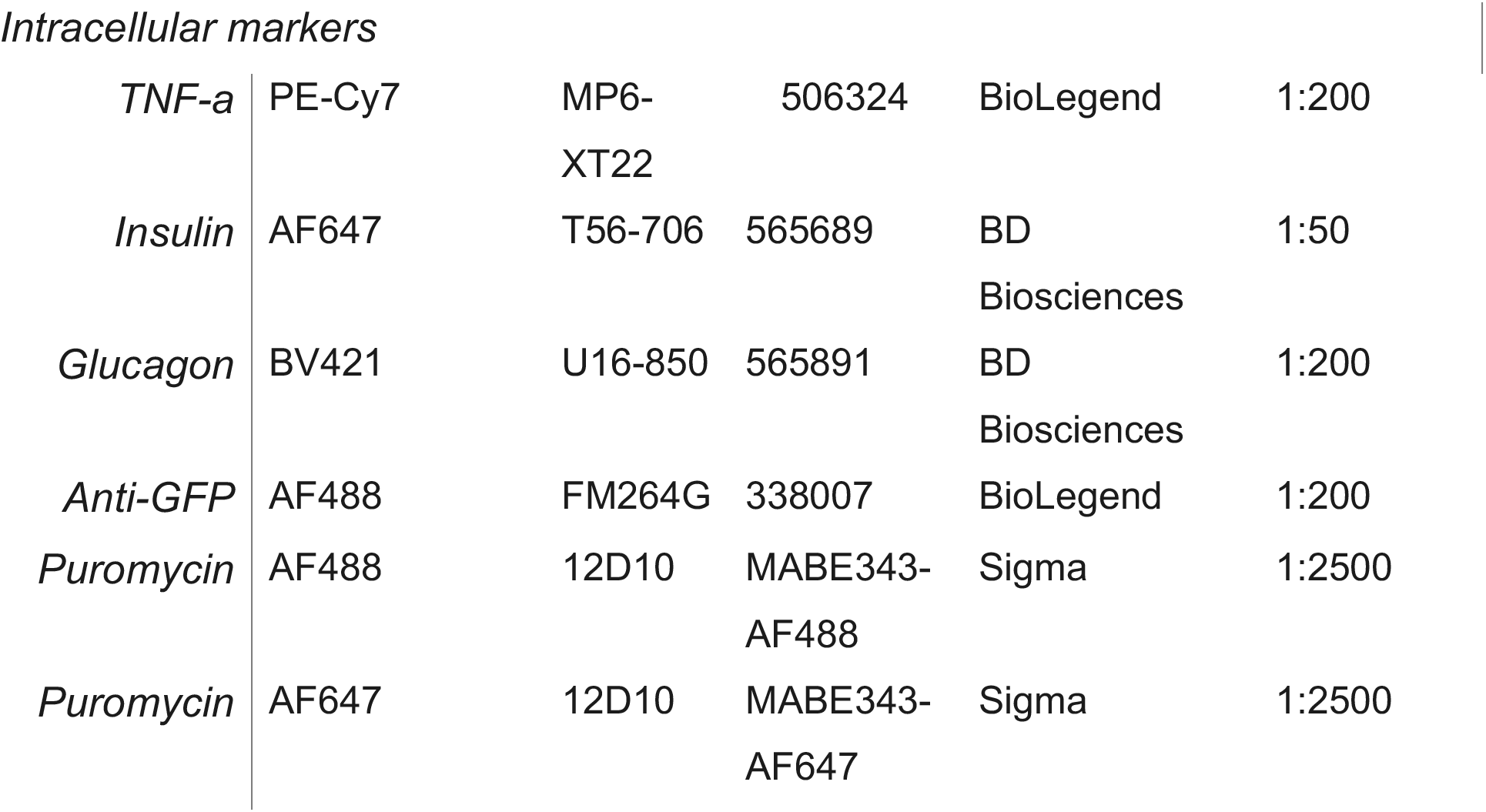
List of antibodies and fluorescent probes used for flow cytometry.

### Cell sorting

Co-cultured GFP MIN6 + BMDMs were gently lifted from the cell culture plate in PBS + 2.5 mM EDTA using cell scrapers. Enrichment of macrophage population for cell sorting was performed using immunomagnetic positive selection for F4/80^+^ cells (EasySep Mouse F4/80 Positive Selection, #100-0659; STEMCELL), according to instructions. Macrophages were then labelled with fixable viability dye (Thermo Fisher #65-0865-18, 1:10000 dilution) and anti-F4/80 (Thermo Fisher #17-4801-82, 1:200 dilution) for 30 min at 4°C in the dark, followed by wash a wash with FACS buffer. Macrophages were transferred to sterile FACS tubes and sorted for live cells, F4/80^+^, and GFP positive or GFP negative (GFP^+^, GFP^-^) into a sterile 15 mL Falcon tube containing 5 mL of 30% FBS in PBS using a BD FACSAria Fusion Cell Sorter (BD) with a 100 µm nozzle. Cells were then spun down, and dry pellets were stored at –80 °C overnight before RNA extraction for Bulk RNA Sequencing, as described below.

### SCENITH

75-100 islets from PhAM^fl^Ins1^Cre^ mice or WT control were transferred into non-tissue culture-treated 48-well plates. Basal condition refers to DMSO control incubation at 37 °C with 5% CO_2_ for 15 min. 100 mM 2-deoxy-d-glucose (2-DG, #D8375; Sigma) was added to 2-DG condition and 2-DG + Oligomycin (DGO) condition and incubated at 37 °C for 5 min. Next, 1.5 µM Oligomycin (Oligo, #O4876; Sigma) was added to Oligo condition, and DGO condition and cells were incubated for at 37 °C for another 10 min. Total incubation time per condition: Basal = 15 min; 2-DG = 15 min, Oligo = 10 min; DGO = 15 min (2-DG for the entire 15 min and Oligo for the final 10 min). 10 µg/mL Puromycin (#P8833; Sigma) was then added to every well, except No Puromycin control, and incubated at 37 °C for 30 min. All samples were then collected into FACS tubes for removal of the supernatant and islet dispersion, as mentioned above. Puromycin fluorescence intensity was assessed in different islet cell types by intracellular flow cytometry staining.

### Fluorescent microscopy

Live cell imaging of GFP MIN6 and co-cultures GFP MIN + BMDMs was performed using a digital inverted benchtop EVOS M5000 fluorescent microscope (Invitrogen). Live cells were imaged in cell culture media at 20x and 40x magnification using transmitted-light (TRANS) and GFP channels.

### Live pancreas imaging

Pancreas from WT and PhAM^fl^Ins1^Cre^ mice were harvested and embedded using 5% agarose in PBS into a cryomold. Blocks of pancreatic tissue embedded in agarose were sliced (200μm) in sterile environment using a vibratome and transferred onto a microscope slide. Tissue slices were kept wet with PBS within the limits of water-repellent A-PAP pen drawing. Tissue slices were blocked using 100μL of Dako Protein Block (Agilent #X0909) containing rat serum (1:50 dilution) and incubated for 10 min with shaking (100 rpm). Slides were washed with PBS and incubated with Fc Block (BD #553141) for 10 min. Antibody staining mix (Table 2) was prepared in Dako Antibody Diluent (Agilent #S3022) and 150μL of staining mix was used per slide. Staining was performed by incubating slides with staining mix in the dark with rotation for 1 h. GFP signal was detected from beta cell mitochondria from PhAM^fl^Ins1^Cre^ mice. Images were acquired using a Leica SP8 in vivo confocal imaging system. Images were analyzed using Leica Application Suite X (LAS X, Leica Microsystmes) Software.

**Table 2.**
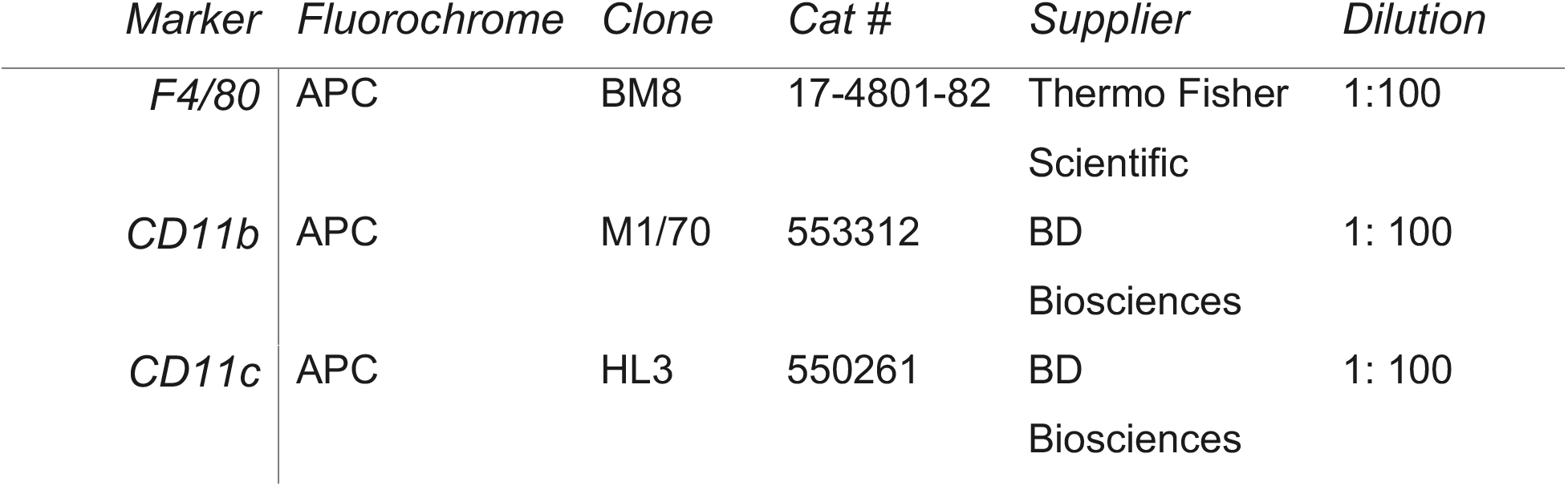
List of antibodies used for confocal microscopy.

### Bulk RNA sequencing

Following cell sorting, RNA was isolated from frozen macrophage pellets using RNeasy Plus Mini Kit (Qiagen #74134), according to manufacturer’s instructions. RNA pellets were submitted for RNA sequencing through the School of Biomedical Engineering (SBME) at the University of British Columbia. RNA sample quality control was performed using the Agilent 2100 Bioanalyzer. Sample preparation was performed following the standard protocol for the Illumina Stranded mRNA preparation. Sequencing was performed using the Illumina NextSeq2000 with Paired End 59bp x 59bp reads. Sequencing data was demultiplexed using Illumina’s BCL Convert app v1.3.0. Adapter trimming was performed by BCL Convert. Read quality assessment, alignment and quantification were performed using the DRAGEN RNA app v4.2.7 on Basespace Sequence Hub with the mouse reference mm10 (annotation Ensembl release 97).

Differential expression analysis was performed with DESeq2 (v1.34.0), with size factors and gene-wise dispersions estimated by the standard procedure and group comparisons assessed by Wald tests. P values were adjusted for multiple testing by the Benjamini–Hochberg method, and genes with an adjusted P < 0.05 were considered differentially expressed. For sample-level visualisation, counts were transformed with the variance-stablishing transformation; overall sample relationships were examined by principal component analysis, and the most significant differentially expressed genes were displayed as per-gene Z-scored heatmaps. KEGG pathway over-representation was assessed separately for the vehicle and cytokine conditions, among genes differentially expressed between GFP^+^ and GFP^-^macrophages, using clusterProfiler (v4.2.2) and *Mus musculus* KEGG gene sets with Benjamini-Hochberg-adjusted P<0.05; enriched pathways are shown as dot plots. Gene identifiers were mapped using the org.Mm.eg.db annotation package. All analyses were conducted in R (v4.1.0).

### Statistical analyses

Data are shown as mean ± SEM. Generation of graphs and statistical analyses were performed using Prism GraphPad version 10. Two-tailed Student’s t test was used when comparing two means. One-way ANOVA with Turkey’s multiple comparison test was applied for more than two groups, and two-way ANOVA was used for two independent variables in two or more groups. Bonferroni’s or Turkey’s post-hoc test was used to compare different sets of means. P values of 0.05 or lower were considered significant.

## Data availability

Further information and requests for resources and reagents should be directed to and will be made available upon reasonable request by the Lead Contacts, Ld Brito Monteiro, RI Klein Geltink, C Bruce Verchere.

All schematics were created using BioRender.

## Acknowledgements

The authors wish to acknowledge Dr. JingSong Wang, Dr. Lisa Xu, BC Children’s Hospital Research Institute (BCCHR) Core Technologies and Services (Vancouver, Canada), for their invaluable assistance with fluorescence imaging and cell sorting. Genomics bioinformatician Bernie Zhao and the School of Biomedical Engineering (SBME) at the University of British Columbia, for RNA sequencing services. Joni Schinkel for imaging editing. The authors also extend their appreciation to the agencies that funded this work: Breakthrough T1D Postdoctoral Fellowship - 3-PDF-2024-1504-A-N (LdBM); Banting Postdoctoral Fellowship - 509775 (ASA).Canadian Institutes of Health Research - AWD-020766 CIHR 2021 (CBV, RKG); CIHR PJ9-198277 (CBV); Investigator award from BC Children’s Hospital Research Institute (CBV); Irving K Barber Chair in Diabetes Research (CBV); Canadian Institutes for Health Research (DT4-179512 to RKG), Natural Sciences and Engineering Research Council of Canada (RGPIN-2020-05390 to RKG), Michael Smith Health Research BC Scholar award (SCH-2022-2767 to RKG).

---

**Supplemental figure 1.**
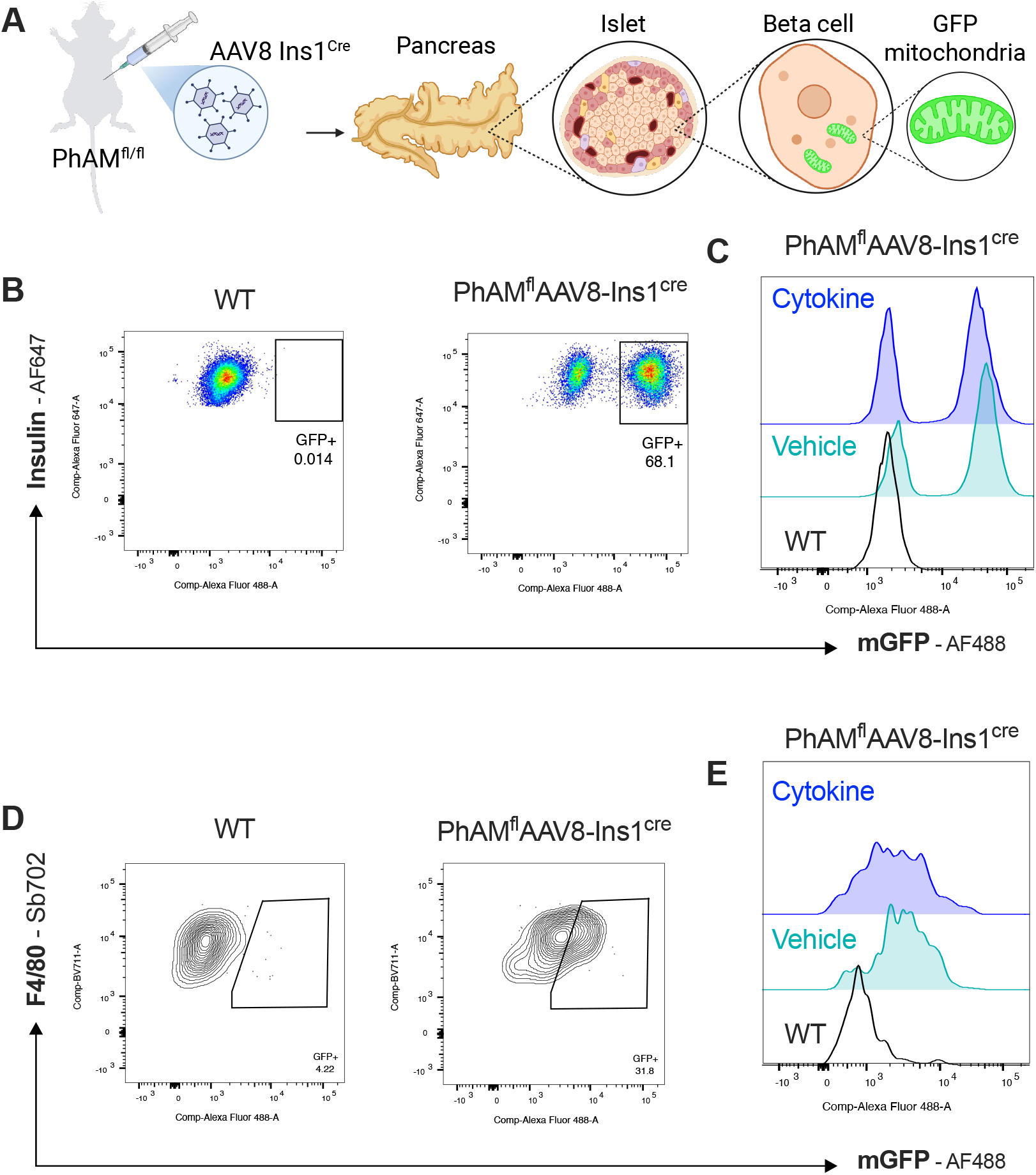
Pancreatic beta cells donate mitochondria to macrophages. Pancreatic islets were collected from Pham^fl^Ins1^Cre^ mice containing a beta cell-specific mitochondrial GFP insert and treated *ex vivo* with either vehicle or pro-inflammatory cytokine cocktail (IL-1β 50 U/mL; TNF-α 1000 U/mL; IFN-γ 1000 U/mL). **A)** Schematics depicting generation of Pham^fl^Ins1^Cre^ mice through injection of AAV8 Ins1^Cre^ directly into the hepatopancreatic duct of PhAM^fl^ mice. **B)** Representative flow plots of mGFP^+^ beta cells (insulin^+^) gated from live cells of whole dispersed islets. **C)** Representative histograms of mGFP^+^ beta cells (insulin^+^) isolated from islets of wild type mice (WT) and Pham^fl^Ins1^Cre^ mice, following treatment with either vehicle or pro-inflammatory cytokines. **D)** Representative flow plots of mGFP^+^ macrophages (F4/80^+^) gated from live cells of whole dispersed islets. **E)** Representative histograms of mGFP^+^ macrophages (F4/80^+^) isolated from islets of wild type mice (WT) and Pham^fl^Ins1^Cre^ mice, following treatment with either vehicle or pro-inflammatory cytokines. Data is shown as mean ± SEM, n = 2 mice.

**Supplemental figure 2.**
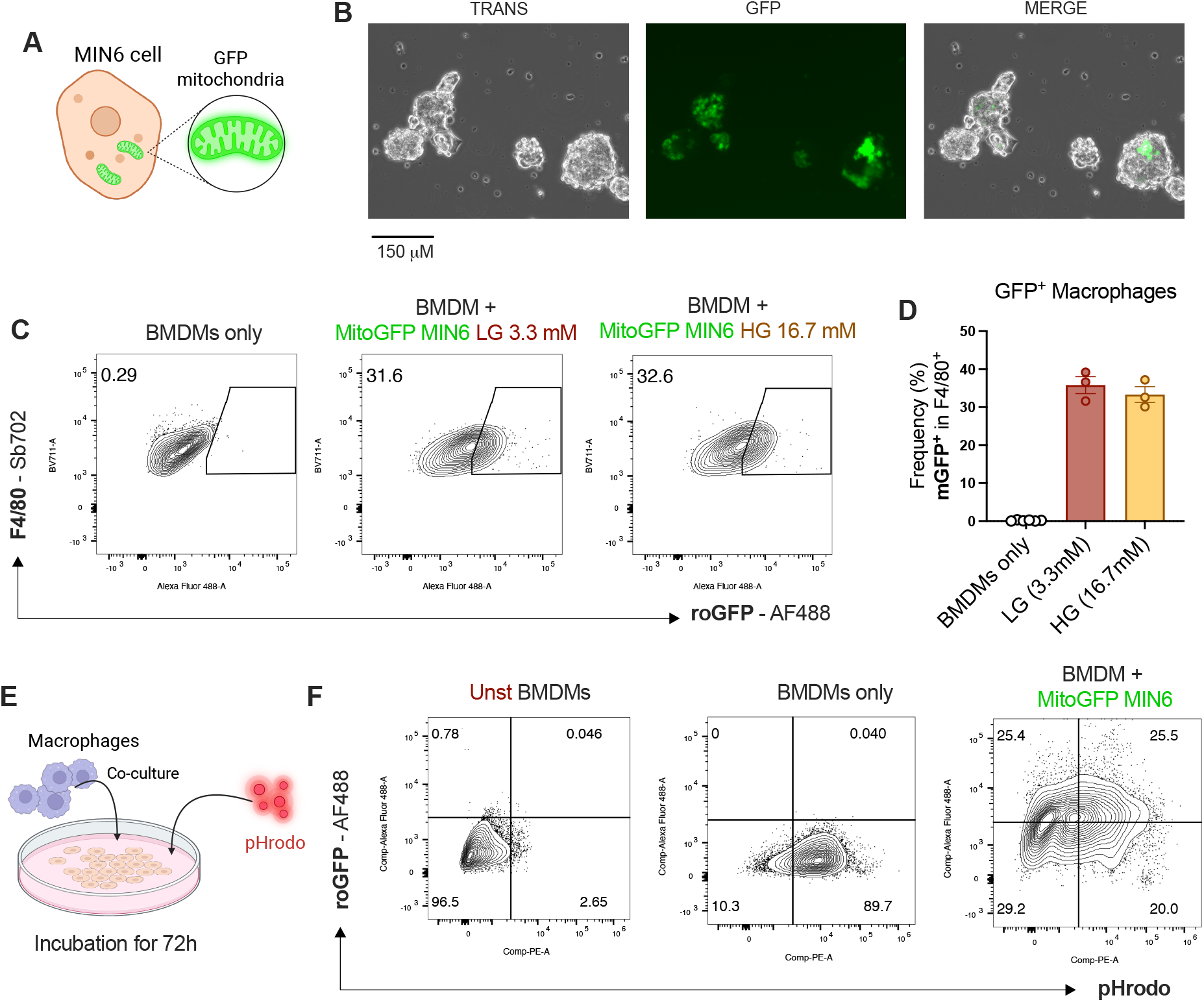
Direct cell culture mediates mitochondrial transfer in vitro in low and high glucose. MIN6 beta cells expressing mitochondrial GFP (MitoGFP) were co-cultured with BMDMs for assessment of mitochondrial transfer events. **A)** Schematics of MIN6 beta cells expressing mitochondrial GFP. **B)** Representative fluorescent microscopy image of MitoGFP MIN6 beta cells. **C)** Representative flow plots of GFP^+^ macrophages (F4/80^+^) directly co-cultured with MitoGFP MIN6 treated with either low glucose 3.3 mM (LG) or high glucose 16.7 mM (HG). **D)** Frequency of GFP^+^ macrophages alone and co-cultured with MitoGFP MIN6 treated with either low glucose 3.3 mM (LG) or high glucose 16.7 mM (HG). **E)** Schematics of long term (72h) direct co-culture of BMDMs + MitoGFP MIN6 beta cells with addition of pHrodo for 30 min for assessment of phagocytosis**. F)** Representative flow plots of pHrodo^+^ / GFP^+^ macrophages (gated in F4/80^+^) co-cultured with or MitoGFP MIN6 for 72h. Data is shown as mean ± SEM, n ≥ 3 co-cultures.

**Supplemental figure 3.**
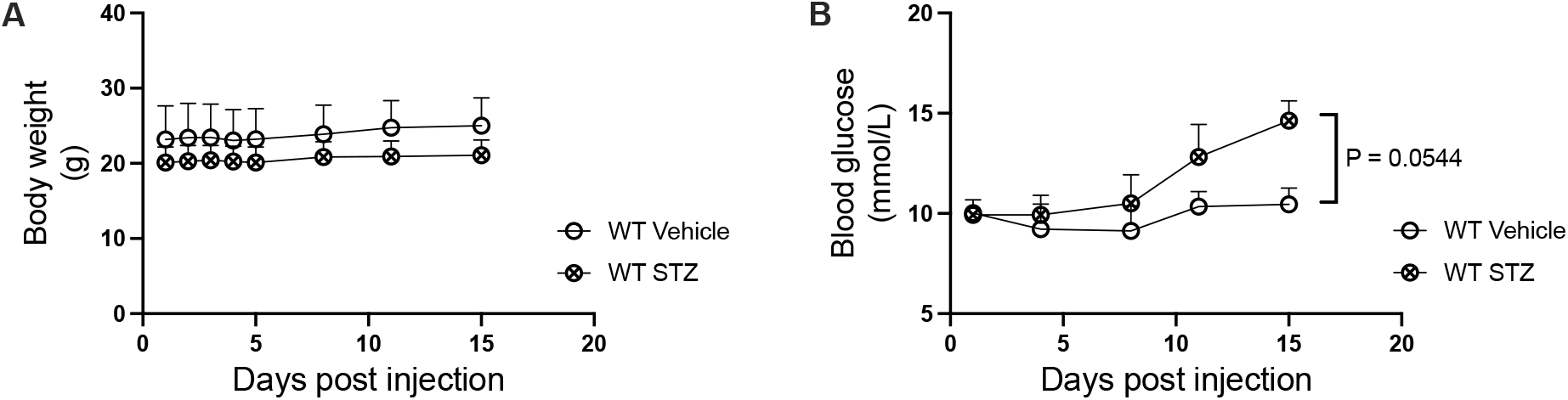
Mouse monitoring post STZ injections. WT and Pham^fl^Ins1^Cre^ mice were treated with 30 mg/kg of STZ for 5 consecutive days and islets were harvested 10 days after the last STZ injection. **A)** Body weight of WT mice treated with vehicle or STZ. **B)** Blood glucose of WT mice treated with vehicle or STZ. n ≥ 3 mice per experiment. P values < 0.05 are considered significant.

**Supplemental figure 4.**
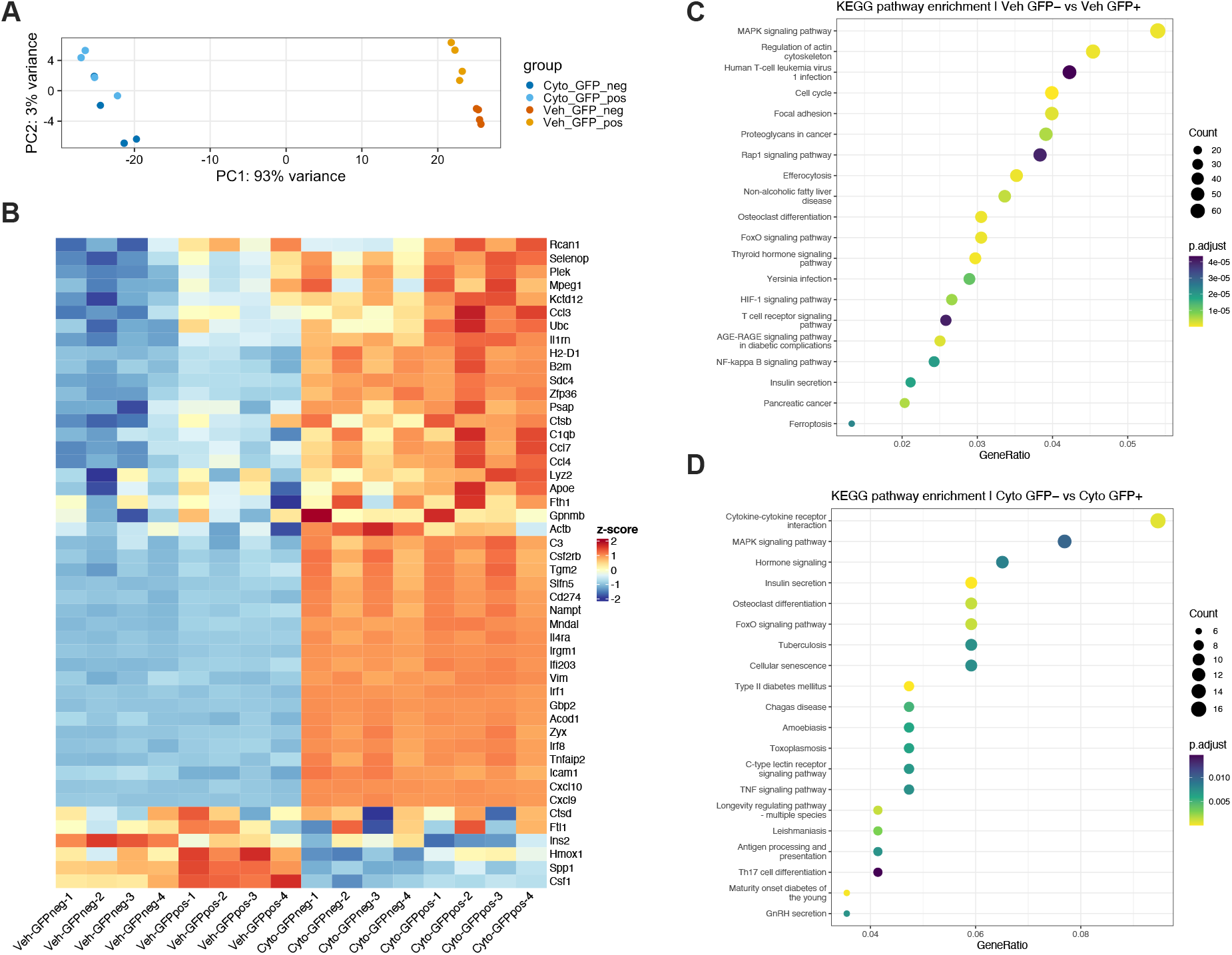
Analysis of differentially expressed genes and pathways in GFP+ and GFP- macrophages. GFP^+^ and GFP^-^ macrophages were sorted and submitted to bulk RNA sequencing. **A)** Principal component analysis of vehicle (Veh)- and cytokine (Cyto)-treated GFP^+^ and GFP^-^ macrophages. **B)** Heatmap of the top 50 differentially expressed genes (ranked by adjusted P value) for vehicle (Veh)- and cytokine (Cyto)-treated GFP^+^ and GFP^-^ macrophages. Rows are genes and columns are individual replicates of the compared groups. **C)** KEGG pathway enrichment of Veh GFP^+^ and GFP^-^ macrophages, and **D)** Cyto GFP^+^ and GFP^-^ macrophages. n=4 per group, P adjusted value <0.05.

## References

1. Carrero, J.A., McCarthy, D.P., Ferris, S.T., Wan, X., Hu, H., Zinselmeyer, B.H., Vomund, A.N., and Unanue, E.R. (2017). Resident macrophages of pancreatic islets have a seminal role in the initiation of autoimmune diabetes of NOD mice. Proc. Natl. Acad. Sci. U. S. A. 114, E10418–E10427. 10.1073/PNAS.1713543114;ISSUE:ISSUE:DOI.

2. Zirpel, H., and Roep, B.O. (2021). Islet-Resident Dendritic Cells and Macrophages in Type 1 Diabetes: In Search of Bigfoot’s Print. Front. Endocrinol. (Lausanne). 12, 666795. 10.3389/FENDO.2021.666795.

3. Mills, E.L., Kelly, B., and O’Neill, L.A.J. (2017). Mitochondria are the powerhouses of immunity. Nature Immunology 2017 18:5 18, 488–498. 10.1038/ni.3704.

4. Weinberg, S.E., Sena, L.A., and Chandel, N.S. (2015). Mitochondria in the regulation of innate and adaptive immunity. Immunity 42, 406–417. 10.1016/j.immuni.2015.02.002.

5. Kelly, B., and O’Neill, L.A.J. (2015). Metabolic reprogramming in macrophages and dendritic cells in innate immunity. Cell Res. 25, 771–784. 10.1038/CR.2015.68.

6. Witcoski Junior, L., de Lima, J.D., Somensi, A.G., de Souza Santos, L.B., Paschoal, G.L., Uada, T.S., Bastos, T.S.B., de Paula, A.G.P., Dos Santos Luz, R.B., Czaikovski, A.P., et al. (2024). Metabolic reprogramming of macrophages in the context of type 2 diabetes. European Journal of Medical Research 2024 29:1 29, 497-. 10.1186/S40001-024-02069-Y.

7. Sharma, R.B., Landa-Galvan, H. V., and Alonso, L.C. (2021). Living Dangerously: Protective and Harmful ER Stress Responses in Pancreatic β-Cells. Diabetes 70, 2431–2443. 10.2337/DBI20-0033.

8. van Tienhoven, R., O’Meally, D., Scott, T.A., Morris, K. V., Williams, J.C., Kaddis, J.S., Zaldumbide, A., and Roep, B.O. (2025). Genetic protection from type 1 diabetes resulting from accelerated insulin mRNA decay. Cell 188, 2407–2416.e9. 10.1016/j.cell.2025.02.018.

9. Bhatti, J.S., Sehrawat, A., Mishra, J., Sidhu, I.S., Navik, U., Khullar, N., Kumar, S., Bhatti, G.K., and Reddy, P.H. (2022). Oxidative stress in the pathophysiology of type 2 diabetes and related complications: Current therapeutics strategies and future perspectives. Free Radic. Biol. Med. 184, 114–134. 10.1016/J.FREERADBIOMED.2022.03.019.

10. Xu, G., Chen, J., Jing, G., and Shalev, A. (2012). Preventing β-Cell Loss and Diabetes With Calcium Channel Blockers. Diabetes 61, 848–856. 10.2337/DB11-0955.

11. Zhou, R., Tardivel, A., Thorens, B., Choi, I., and Tschopp, J. (2010). Thioredoxin-interacting protein links oxidative stress to inflammasome activation. Nat. Immunol. 11, 136–140. 10.1038/NI.1831.

12. Forlenza, G.P., McVean, J., Beck, R.W., Bauza, C., Bailey, R., Buckingham, B., DiMeglio, L.A., Sherr, J.L., Clements, M., Neyman, A., et al. (2023). Effect of Verapamil on Pancreatic Beta Cell Function in Newly Diagnosed Pediatric Type 1 Diabetes: A Randomized Clinical Trial. JAMA 329, 990–999. 10.1001/JAMA.2023.2064.

13. Darwish, R., Alcibahy, Y., Abu-Sharia, G., and Butler, A.E. (2025). β-Cell Mitochondrial Dysfunction: Underlying Mechanisms and Potential Therapeutic Strategies. Cells 2025, Vol. 14, Page 1861 14, 1861. 10.3390/CELLS14231861.

14. Lytrivi, M., Tong, Y., Virgilio, E., Yi, X., and Cnop, M. (2025). Diabetes mellitus and the key role of endoplasmic reticulum stress in pancreatic β cells. Nature Reviews Endocrinology 2025 21:9 21, 546–563. 10.1038/s41574-025-01129-5.

15. Terasaki, A., Bhatnagar, K., Weiner, A.T., Tan, Y., Szeifert, V., Huang, H.-L., Wiggers, L., Rodrigues, V., Rada, C.C., Shankar, V., et al. (2026). Mitochondrial transfer from immune to tumor cells enables lymph node metastasis. Cell Metab. 38, 388–398.e7. 10.1016/j.cmet.2025.12.014.

16. Saha, T., Dash, C., Jayabalan, R., Khiste, S., Kulkarni, A., Kurmi, K., Mondal, J., Majumder, P.K., Bardia, A., Jang, H.L., et al. (2021). Intercellular nanotubes mediate mitochondrial trafficking between cancer and immune cells. Nature Nanotechnology 2021 17:1 17, 98–106. 10.1038/s41565-021-01000-4.

17. Xu, J., Li, Y., Novak, C., Lee, M., Yan, Z., Bang, S., McGinnis, A., Chandra, S., Zhang, V., He, W., et al. (2026). Mitochondrial transfer from glia to neurons protects against peripheral neuropathy. Nature 2026 650:8103 650, 951–960. 10.1038/s41586-025-09896-x.

18. Brestoff, J.R., Wilen, C.B., Moley, J.R., Li, Y., Zou, W., Malvin, N.P., Rowen, M.N., Saunders, B.T., Ma, H., Mack, M.R., et al. (2021). Intercellular Mitochondria Transfer to Macrophages Regulates White Adipose Tissue Homeostasis and Is Impaired in Obesity. Cell Metab. 33, 270–282.e8. 10.1016/j.cmet.2020.11.008.

19. Baldwin, J.G., Heuser-Loy, C., Saha, T., Schelker, R., Slavkovic-Lukic, D., Strieder, N., Hernandez-Lopez, I., Rana, N., Barden, M., Mastrogiovanni, F., et al. (2024). Intercellular nanotube-mediated mitochondrial transfer enhances T cell metabolic fitness and antitumor efficacy. Cell 187, 6614–6630.e21. 10.1016/j.cell.2024.08.029.

20. Cosentino, C., and Regazzi, R. (2021). Crosstalk between Macrophages and Pancreatic β-Cells in Islet Development, Homeostasis and Disease. Int. J. Mol. Sci. 22, 1765. 10.3390/IJMS22041765.

21. Pham, A.H., Mccaffery, J.M., and Chan, D.C. (2012). Mouse lines with photo-activatable mitochondria to study mitochondrial dynamics. genesis 50, 833–843. 10.1002/DVG.22050.

22. Ramzy, A., Tudurí, E., Glavas, M.M., Baker, R.K., Mojibian, M., Fox, J.K., O’Dwyer, S.M., Dai, D., Hu, X., Denroche, H.C., et al. (2020). AAV8 Ins1-Cre can produce efficient β-cell recombination but requires consideration of off-target effects. Scientific Reports 2020 10:1 10, 10518-. 10.1038/s41598-020-67136-w.

23. Smith, K.A., Waypa, G.B., and Schumacker, P.T. (2017). Redox signaling during hypoxia in mammalian cells. Redox Biol. 13, 228–234. 10.1016/J.REDOX.2017.05.020.

24. Waypa, G.B., Marks, J.D., Guzy, R., Mungai, P.T., Schriewer, J., Dokic, D., and Schumacker, P.T. (2010). Hypoxia triggers subcellular compartmental redox signaling in vascular smooth muscle cells. Circ. Res. 106, 526–535. 10.1161/CIRCRESAHA.109.206334;REQUESTEDJOURNAL:JOURNAL:RES;CTYPE:STRING:JOURNAL.

25. Fan, A.C., Thota, R.R., Serwas, N., Vykunta, V.S., Marchuk, K., Ruhland, M.K., Liu, L., Johnson, G., Edwards, A., and Krummel, M.F. (2026). Submicrometre sampling of living cells by macrophages. Nature 2026 654:8118 654, 495–503. 10.1038/s41586-026-10435-5.

26. Ying, W., Lee, Y.S., Dong, Y., Seidman, J.S., Yang, M., Isaac, R., Seo, J.B., Yang, B.H., Wollam, J., Riopel, M., et al. (2019). Expansion of Islet-Resident Macrophages Leads to Inflammation Affecting β Cell Proliferation and Function in Obesity. Cell Metab. 29, 457–474.e5. 10.1016/j.cmet.2018.12.003.

27. Argüello, R.J., Combes, A.J., Char, R., Gigan, J.P., Baaziz, A.I., Bousiquot, E., Camosseto, V., Samad, B., Tsui, J., Yan, P., et al. (2020). SCENITH: A Flow Cytometry-Based Method to Functionally Profile Energy Metabolism with Single-Cell Resolution. Cell Metab. 32, 1063–1075.e7. 10.1016/j.cmet.2020.11.007.

28. de Brito Monteiro, L., Archambault, A.S., Soukhatcheva, G., Dai, D., Velghe, J., Chen, Y.C., Verchere, C.B., and Klein Geltink, R.I. (2025). Assessment of protein synthesis rate enables metabolic profiling of resident-immune cells of the islets of Langerhans. Front. Immunol. 16, 1662986. 10.3389/FIMMU.2025.1662986/FULL.

29. Nackiewicz, D., Dan, M., Speck, M., Chow, S.Z., Chen, Y.C., Pospisilik, J.A., Verchere, C.B., and Ehses, J.A. (2020). Islet Macrophages Shift to a Reparative State following Pancreatic Beta-Cell Death and Are a Major Source of Islet Insulin-like Growth Factor-1. iScience 23, 100775. 10.1016/J.ISCI.2019.100775.

30. Song, R., Huang, C., Ma, Y., Zhang, Z., Liu, Y., Chen, B., Zhang, X., Hao, S., Huang, H., Ashrafizadeh, M., et al. (2026). Cytoskeletal remodeling promotes tunneling nanotube formation and drives cardiac resident cell mitochondrial transfer in sepsis. Sci. Adv. 12, eadz3266. 10.1126/SCIADV.ADZ3266;PAGEGROUP:STRING:PUBLICATION.

31. Wei, Z., and Liu, H.T. (2002). MAPK signal pathways in the regulation of cell proliferation in mammalian cells. Cell Research 2002 12:1 12, 9–18. 10.1038/sj.cr.7290105.

32. Stulpinas, A., Uzusienis, T., Imbrasaite, A., Krestnikova, N., Unguryte, A., and Kalvelyte, A. V. (2021). Cell-cell and cell-substratum contacts in the regulation of MAPK and Akt signalling: Importance in therapy, biopharmacy and bioproduction. Cell. Signal. 84, 110034. 10.1016/J.CELLSIG.2021.110034.

33. Faust, D., Dolado, I., Cuadrado, A., Oesch, F., Weiss, C., Nebreda, A.R., and Dietrich, C. (2005). p38α MAPK is required for contact inhibition. Oncogene 24, 7941–7945. 10.1038/SJ.ONC.1208948;KWRD.

34. Bukoreshtliev, N. V., Wang, X., Hodneland, E., Gurke, S., Barroso, J.F.V., and Gerdes, H.H. (2009). Selective block of tunneling nanotube (TNT) formation inhibits intercellular organelle transfer between PC12 cells. FEBS Lett. 583, 1481–1488. 10.1016/J.FEBSLET.2009.03.065;SUBPAGE:STRING:ABSTRACT;WGROUP:STRING:PUBLICATION.

35. Brož, A., Volfová, T., Javorská, Ž., Rösel, D., Brábek, J., and Perlíková, P. (2026). The potential of cytochalasins and other cytochalasans as drugs targeting cancer cell invasiveness and metastasis. Eur. J. Med. Chem. 304, 118502. 10.1016/J.EJMECH.2025.118502.

36. Spees, J.L., Olson, S.D., Whitney, M.J., and Prockop, D.J. (2006). Mitochondrial transfer between cells can rescue aerobic respiration. Proc. Natl. Acad. Sci. U. S. A. 103, 1283–1288. 10.1073/PNAS.0510511103;WEBSITE:WEBSITE:PNAS-SITE;ISSUE:ISSUE:DOI.

37. Islam, M.N., Das, S.R., Emin, M.T., Wei, M., Sun, L., Westphalen, K., Rowlands, D.J., Quadri, S.K., Bhattacharya, S., and Bhattacharya, J. (2012). Mitochondrial transfer from bone-marrow–derived stromal cells to pulmonary alveoli protects against acute lung injury. Nature Medicine 2012 18:5 18, 759–765. 10.1038/nm.2736.

38. Liu, D., Gao, Y., Liu, J., Huang, Y., Yin, J., Feng, Y., Shi, L., Meloni, B.P., Zhang, C., Zheng, M., et al. (2021). Intercellular mitochondrial transfer as a means of tissue revitalization. Signal Transduction and Targeted Therapy 2021 6:1 6, 65-. 10.1038/s41392-020-00440-z.

39. Rodriguez, A.M., Nakhle, J., Griessinger, E., and Vignais, M.L. (2018). Intercellular mitochondria trafficking highlighting the dual role of mesenchymal stem cells as both sensors and rescuers of tissue injury. Cell Cycle 17, 712. 10.1080/15384101.2018.1445906.

40. Cruz, A.F., Rohban, R., and Esni, F. (2020). Macrophages in the pancreas: Villains by circumstances, not necessarily by actions. Immun. Inflamm. Dis. 8, 807–824. 10.1002/IID3.345;SUBPAGE:STRING:FRAMEDPDF;PAGE:STRING:ARTICLE/CHAPTER.

41. Caronni, N., La Terza, F., Vittoria, F.M., Barbiera, G., Mezzanzanica, L., Cuzzola, V., Barresi, S., Pellegatta, M., Canevazzi, P., Dunsmore, G., et al. (2023). IL-1β+ macrophages fuel pathogenic inflammation in pancreatic cancer. Nature 2023 623:7986 623, 415–422. 10.1038/s41586-023-06685-2.

42. Nackiewicz, D., Dan, M., He, W., Kim, R., Salmi, A., Rütti, S., Westwell-Roper, C., Cunningham, A., Speck, M., Schuster-Klein, C., et al. (2014). TLR2/6 and TLR4-activated macrophages contribute to islet inflammation and impair beta cell insulin gene expression via IL-1 and IL-6. Diabetologia 2014 57:8 57, 1645–1654. 10.1007/S00125-014-3249-1.

43. Ying, W., Fu, W., Lee, Y.S., and Olefsky, J.M. (2020). The role of macrophages in obesity-associated islet inflammation and β-cell abnormalities. Nat. Rev. Endocrinol. 16, 81–90. 10.1038/S41574-019-0286-3;SUBJMETA.

44. Ashcroft, F.M., Lloyd, M., and Haythorne, E.A. (2023). Glucokinase activity in diabetes: too much of a good thing? Trends in Endocrinology and Metabolism 34, 119–130. 10.1016/j.tem.2022.12.007.

45. O’Neill, L.A., Kishton, R.J., and Rathmell, J. (2016). A guide to immunometabolism for immunologists. Nat Rev Immunol 16, 553–565. 10.1038/nri.2016.70.

46. den Bossche, J., O’Neill, L.A., and Menon, D. (2017). Macrophage Immunometabolism: Where Are We (Going)? Trends Immunol 38, 395–406. 10.1016/j.it.2017.03.001.

47. Klinkhammer, C., Dohle, C., and Gleichmann, H. (1989). T Cell-Dependent Class II Major Histocompatibility Complex Antigen Expression in vivo Induced by the Diabotegen Streptozotocin. Immunobiology 180, 1–11. 10.1016/S0171-2985(89)80025-7.

48. Zou, H., Xie, L., Hu, J., Zhang, R., Wang, Y., Xu, A., Zhou, Z., Xiao, X., and Xiao, Y. (2025). Fatty Acid Binding Protein 4 Regulates the Antigen-Presenting Function of Dendritic Cells Resulting in T Cell Priming in Streptozotocin-Induced Type 1 Diabetes Mice. J. Diabetes 17, e70123. 10.1111/1753-0407.70123;PAGEGROUP:STRING:PUBLICATION.

49. Scheiblich, H., Dansokho, C., Mercan, D., Schmidt, S. V., Bousset, L., Wischhof, L., Eikens, F., Odainic, A., Spitzer, J., Griep, A., et al. (2021). Microglia jointly degrade fibrillar alpha-synuclein cargo by distribution through tunneling nanotubes. Cell 184, 5089–5106.e21. 10.1016/J.CELL.2021.09.007.

50. Alarcon-Martinez, L., Villafranca-Baughman, D., Quintero, H., Kacerovsky, J.B., Dotigny, F., Murai, K.K., Prat, A., Drapeau, P., and Di Polo, A. (2020). Interpericyte tunnelling nanotubes regulate neurovascular coupling. Nature 2020 585:7823 585, 91–95. 10.1038/s41586-020-2589-x.

51. Sun, C., Liu, X., Wang, B., Wang, Z., Liu, Y., Di, C., Si, J., Li, H., Wu, Q., Xu, D., et al. (2019). Endocytosis-mediated mitochondrial transplantation: Transferring normal human astrocytic mitochondria into glioma cells rescues aerobic respiration and enhances radiosensitivity. Theranostics 9, 3595–3607. 10.7150/THNO.33100.

52. Wang, Y., Yu, H.Y., Yi, Z.J., Qi, L.Y., Yang, J.S., Xie, H.X., Zhao, M., Liu, N.H., Chen, J.Q., Zhou, T.J., et al. (2025). Super mitochondria-enriched extracellular vesicles enable enhanced mitochondria transfer. Nature Communications 2025 16:1 16, 9448-. 10.1038/s41467-025-64486-9.

53. Hayakawa, K., Esposito, E., Wang, X., Terasaki, Y., Liu, Y., Xing, C., Ji, X., and Lo, E.H. (2016). Transfer of mitochondria from astrocytes to neurons after stroke. Nature 2016 535:7613 535, 551–555. 10.1038/nature18928.

54. Liu, D., Gao, Y., Liu, J., Huang, Y., Yin, J., Feng, Y., Shi, L., Meloni, B.P., Zhang, C., Zheng, M., et al. (2021). Intercellular mitochondrial transfer as a means of tissue revitalization. Signal Transduction and Targeted Therapy 2021 6:1 6, 1–18. 10.1038/s41392-020-00440-z.

55. Maeda, A., and Fadeel, B. (2014). Mitochondria released by cells undergoing TNF-α-induced necroptosis act as danger signals. Cell Death & Disease 2014 5:7 5, e1312–e1312. 10.1038/cddis.2014.277.

56. Ufer, F., Vargas, P., Engler, J.B., Tintelnot, J., Schattling, B., Winkler, H., Bauer, S., Kursawe, N., Willing, A., Keminer, O., et al. (2016). Arc/Arg3.1 governs inflammatory dendritic cell migration from the skin and thereby controls T cell activation. Sci. Immunol. 1. 10.1126/SCIIMMUNOL.AAF8665.

57. Song, R., Huang, C., Ma, Y., Zhang, Z., Liu, Y., Chen, B., Zhang, X., Hao, S., Huang, H., Ashrafizadeh, M., et al. (2026). Cytoskeletal remodeling promotes tunneling nanotube formation and drives cardiac resident cell mitochondrial transfer in sepsis. Science Advances 12, 1–18. 10.1126/SCIADV.ADZ3266;PAGEGROUP:STRING:PUBLICATION.

58. Soukhatcheva, G., Stanley, L., Dai, L., Komba, M., Andriiets, V., Johnson, J.D., Verchere, C.B., and Chen, Y.C. (2026). Manipulation of Gene Expression in Mouse Pancreas via Intraductal Delivery of Adeno-Associated Viral Vectors. Bio. Protoc. 16. 10.21769/BIOPROTOC.5657.

59. Ramzy, A., Tudurí, E., Glavas, M.M., Baker, R.K., Mojibian, M., Fox, J.K., O’Dwyer, S.M., Dai, D., Hu, X., Denroche, H.C., et al. (2020). AAV8 Ins1-Cre can produce efficient β-cell recombination but requires consideration of off-target effects. Scientific Reports 2020 10:1 10, 10518-. 10.1038/s41598-020-67136-w.

60. Smith, N., F Spigelman, A., Lin, H., and E Macdonald, P. (2018). Mouse Pancreatic Islet Isolation v1. 10.17504/PROTOCOLS.IO.SQAEDSE.

